# Spatial coordination of filamentous enzymes establishes a redox-balancing metabolon

**DOI:** 10.64898/2026.11.13.766394

**Authors:** Nemanja Kutlešić, Sean Krivitsky, Jed de Ruiter-Swain, Yijian Qiu, Sangjun Yoon, Wen-Hsuan Yang, Sara Pawlak, Paolo Cifani, Tse-Luen Wee, Michael J. Lukey

## Abstract

Nutrient scarcity in tumors prompts metabolic adaptations in cancer cells. Here, we identify that glutamine deprivation, a common feature of the tumor microenvironment, induces formation of a proline biosynthesis metabolon to selectively maintain intracellular proline levels. Mechanistically, filament-forming enzymes glutaminase (GLS) and pyrroline-5-carboxylate synthase (P5CS) co-assemble with the downstream reductases PYCR1 and PYCR2 into a physically associated enzyme complex within a spatially segregated mitochondrial subset. This assembly is dynamically regulated by the pathway intermediate glutamate and is membrane-anchored via ATP synthase. Disruption of enzyme spatial coordination compromises pathway activity, while the sustained proline biosynthesis is repurposed to support redox homeostasis under limited glutamine. We reveal nutrient-responsive filamentation and spatial coordination of mitochondrial enzymes as a mechanism of rapid metabolic adaptation to the fluctuating cellular environment.

## Main

Cancer cells frequently encounter nutrient scarcity within solid tumors as a result of uneven vascular perfusion and intense competition for available resources^1–3^. Survival under these conditions requires metabolic adaptations that preserve essential cellular functions when supplies of many biosynthetic precursors and metabolic fuels are diminished^1,4^. Because individual nutrients typically support multiple metabolic processes, the capacity to favor certain pathways over others may become particularly important when a shared substrate is limiting. Exemplifying this challenge, glutamine serves as a major carbon and nitrogen source for several metabolic pathways, yet its availability can become severely restricted within the tumor microenvironment (TME)^1,4,5^.

The activity of a metabolic pathway depends not only on substrate availability and the expression levels and catalytic properties of its constituent enzymes, but also on how these enzymes are spatially organized within the cell^6–9^. Enzymes catalyzing sequential reactions can assemble into higher-order complexes, termed metabolons, which bring pathway components into close proximity to support coordinated activity^6,8,10,11^. Metabolons can assemble and disassemble in response to changes in metabolite availability, enabling rapid reorganization of metabolic pathways without corresponding changes in enzyme abundance^6,12^. Another form of higher-order molecular organization is metabolic enzyme filamentation, in which multiple units of the same enzyme reversibly polymerize into extended structures^13–16^. Filament assembly is often responsive to changes in substrate and/or product availability and can modulate catalytic activity by stabilizing specific conformations of the enzyme^17–19^. However, whether filament-forming enzymes participate in physically associated multienzyme complexes that sustain pathway function and how they may influence the allocation of limiting substrates among competing metabolic processes remains poorly understood.

### Glutamine deprivation induces coordinated redistribution of proline biosynthesis enzymes

To investigate whether specific metabolic processes are preferentially maintained under conditions of limited glutamine availability, we analyzed the abundance of glutamine-derived metabolites in glutamine-dependent human triple-negative breast cancer (TNBC) cells after 24 h of culture in glutamine-replete (4 mM) or glutamine-depleted (25 µM) conditions. As expected, glutamine limitation caused a pronounced decrease in the abundance of most glutamine-derived metabolites, including glutamate and α-ketoglutarate (Fig. 1A). However, intracellular proline levels were selectively maintained (Fig. 1A). This preservation was not attributable to increased synthesis from arginine, as stable-isotope tracing revealed little incorporation of arginine-derived carbon into proline under either condition, despite substantial labeling of ornithine (Supplementary Fig. 1A).

**Figure 1.**
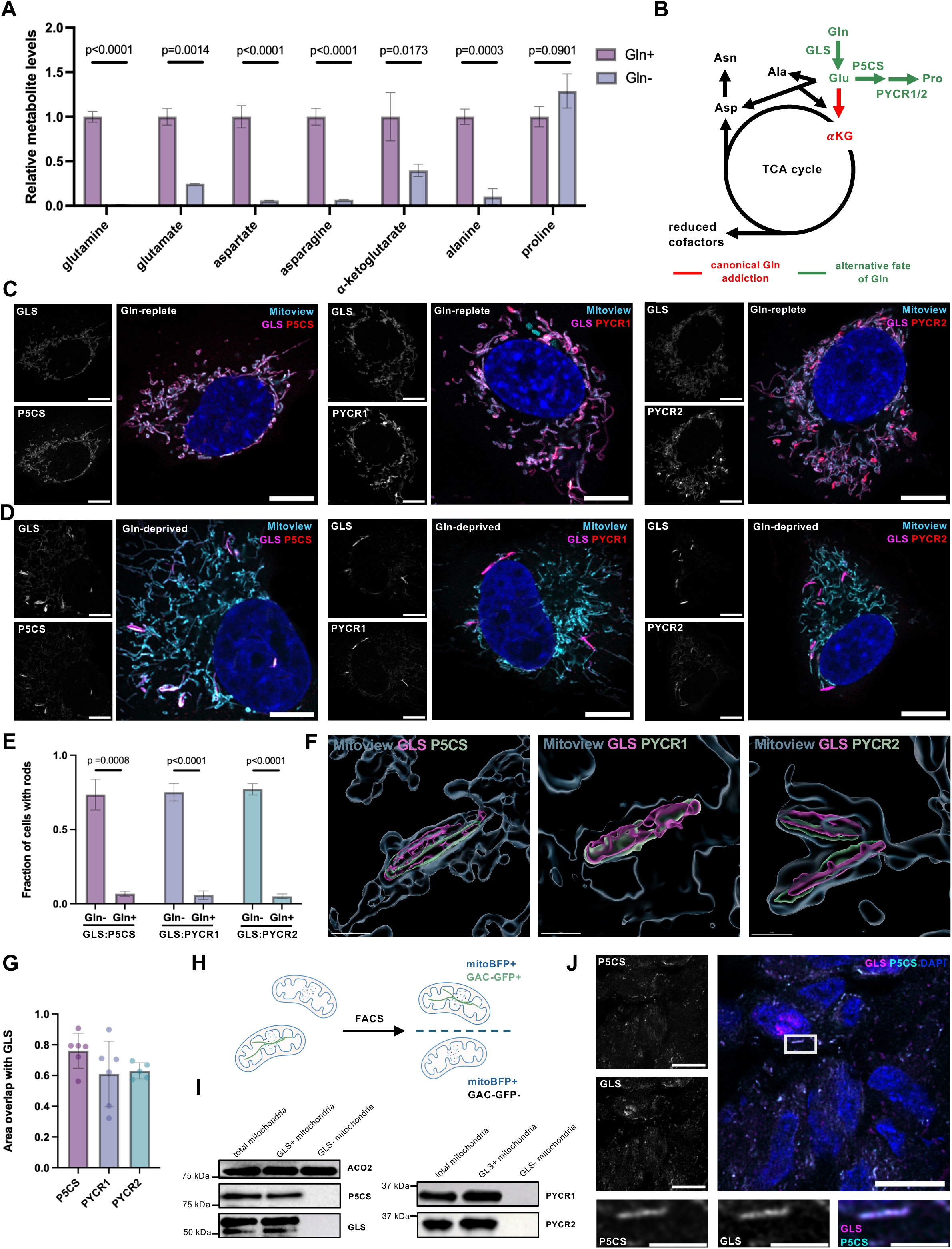
Glutamine deprivation drives coordinated segregation of proline biosynthesis enzymes into a distinct mitochondrial subpopulation. **(A)** Relative metabolite levels in MDA- MB-231 cells under glutamine-replete (4 mM) and glutamine-deprived (25 *μ*M) conditions, normalized to glutamine-replete conditions. Data from *N=*3 independent replicates. Statistical significance determined on log2-transformed data using the two-tailed Student’s t-test with Welch’s correction. **(B)** Schematic of downstream metabolic fates of glutamine in glutamine-dependent cancer cells, with proline maintenance highlighted in green. **(C)** Representative immunofluorescence images of GLS, P5CS, PYCR1 and PYCR2 distribution within the mitochondrial network under glutamine-replete conditions (4 mM) in Hs 578T cells. Cells grown in DMEM. Scale bar 10 *μ*m. (D) Representative immunofluorescence images of GLS, P5CS, PYCR1 and PYCR2 distribution within the mitochondrial network under glutamine-deprived conditions (25 *μ*M) in Hs 578T cells. Cells grown in DMEM. Scale bar 10 *μ*m. **(E)** Quantified fractions of cells with cellular rods of all indicated pairs of enzymes in the proline biosynthesis pathway, under glutamine-deprived and glutamine-replete conditions. Data from *N=*3 independent replicates. Statistical significance determined using the two-tailed Student’s t-test. **(F)** 3D reconstructions of GLS-, P5CS-, PYCR1-, and PYCR2-enriched mitochondrial rods under glutamine deprivation (25 *μ*M). Scale bar 1 *μ*m. **(G)** Quantified area overlap of GLS with total signal for each of the proline biosynthesis enzymes in subpopulations under glutamine deprivation. **(H)** Schematic of FACS-based assay for separation of GLS-containing and GLS-devoid mitochondrial subpopulations. **(I)** Western blots on segregated mitochondrial subpopulations. **(J)** Representative immunofluorescence image of a human breast tumor slice co-stained for GLS and P5CS, showing rod morphology for both enzymes. Scale bar 10 *μ*m. Insets show an individual rod, scale bar 2 *μ*m.

The selective preservation of proline levels prompted us to examine whether glutamine deprivation alters either the expression levels or intracellular organization of enzymes involved in its biosynthesis (Fig. 1B). Under glutamine-replete conditions, glutaminase (GLS), which catalyzes the initial hydrolysis of glutamine to glutamate and ammonium^4,5^, was distributed throughout the mitochondrial network (Supplementary Fig. 1B). Upon glutamine deprivation, however, GLS became concentrated within a distinct subpopulation of elongated mitochondria, producing prominent GLS-enriched rods (Supplementary Fig. 1C-E). Such structures have previously been linked to reversible polymerization into enzyme filaments^20–23^. Super-resolution Airyscan microscopy resolved multiple elongated GLS-positive foci within these mitochondria, rather than a single continuous structure (Supplementary Fig. 1F). This reorganization occurred without a detectable change in GLS protein abundance (Supplementary Fig. 1G). In contrast, GLS remained broadly distributed during glucose or serum deprivation, as well as during culture in galactose media, to increase reliance on oxidative phosphorylation, supporting the specificity of this response to glutamine deprivation (Supplementary Fig. 2A-E).

Downstream of GLS, pyrroline-5-carboxylate synthase (P5CS) converts glutamate to P5C, which is subsequently reduced to proline by P5C reductases 1 and 2 (PYCR1/2) (Fig. 1B). Strikingly, P5CS, PYCR1, and PYCR2 also shifted from a pan-mitochondrial distribution and co-segregated with GLS following glutamine deprivation (Fig. 1C -E), forming elongated rod-shaped structures. Three-dimensional reconstructions revealed spatial overlap between GLS and each of the downstream enzymes under these conditions (Fig. 1F, G, Supplementary Movies 1-3). The same pattern of enzyme reorganization was observed in multiple glutamine-dependent human breast cancer cell lines (Supplementary Fig. 3A-F). In line with the selective maintenance of proline, co-segregation was specific to the proline biosynthesis pathway, with other mitochondrial enzymes involved in glutamine metabolism, such as aspartate aminotransferase 2 (GOT2) and aconitase 2 (ACO2), retaining their pan-mitochondrial distribution under the same conditions (Supplementary Fig. 4A, B). The redistribution of GLS, P5CS, PYCR1, and PYCR2 occurred without detectable changes in their protein abundance (Supplementary Fig. 4C).

To corroborate the co-segregation of GLS with downstream proline biosynthesis enzymes, we developed a fluorescence-activated mitochondrial sorting strategy based on GFP-tagged GLS. We stably expressed GFP-tagged GLS in TNBC cells in which the endogenous *GLS* gene had been knocked out using CRISPR/Cas9. The GLS-GFP fusion protein underwent the characteristic redistribution into a distinct mitochondrial subpopulation upon glutamine deprivation (Supplementary Fig. 4D). Mitochondria isolated from glutamine-deprived cells expressing the tagged GLS along with a mitochondria-targeted fluorescent marker (mBFP2) were then sorted into GLS-positive and GLS-negative populations (Fig. 1H). Consistent with our microscopy observations, P5CS, PYCR1 and PYCR2 were highly enriched in the GLS-positive mitochondrial fraction relative to the GLS-negative fraction (Fig. 1I). Together, these findings demonstrate that glutamine deprivation induces the selective co-segregation of glutamine-to-proline biosynthesis enzymes, giving rise to a distinct mitochondrial subpopulation enriched in these pathway components.

To determine whether this spatial organization also occurs in human breast cancer, we examined the distribution of GLS and P5CS in specimens of invasive ductal carcinoma. Pronounced co-enrichment of GLS and P5CS within elongated rod-shaped structures was evident (Fig. 1J, Supplementary Fig. 4E), supporting the pathophysiological relevance of our cell culture findings.

### Glutamate availability regulates GLS and P5CS filamentation and co-segregation

The induction of enzyme co-segregation upon glutamine withdrawal raises the question of whether this spatial organization is regulated by glutamine itself or by a downstream metabolite. Because glutamate is both the product of GLS and the substrate of P5CS, we tested whether its availability influences enzyme redistribution during glutamine limitation. Remarkably, glutamate supplementation completely prevented the co-segregation of GLS and P5CS into a distinct mitochondrial subpopulation during glutamine restriction, although smaller punctate structures of P5CS remained detectable in some cells (Fig. 2A). A similar effect was observed for both PYCR1 and PYCR2, which also retained their pan-mitochondrial distribution when glutamine-deficient medium was supplemented with glutamate (Fig. 2B-D). Taken together, these results identify a role for the pathway intermediate glutamate in controlling the coordinated redistribution of proline biosynthesis enzymes.

**Figure 2.**
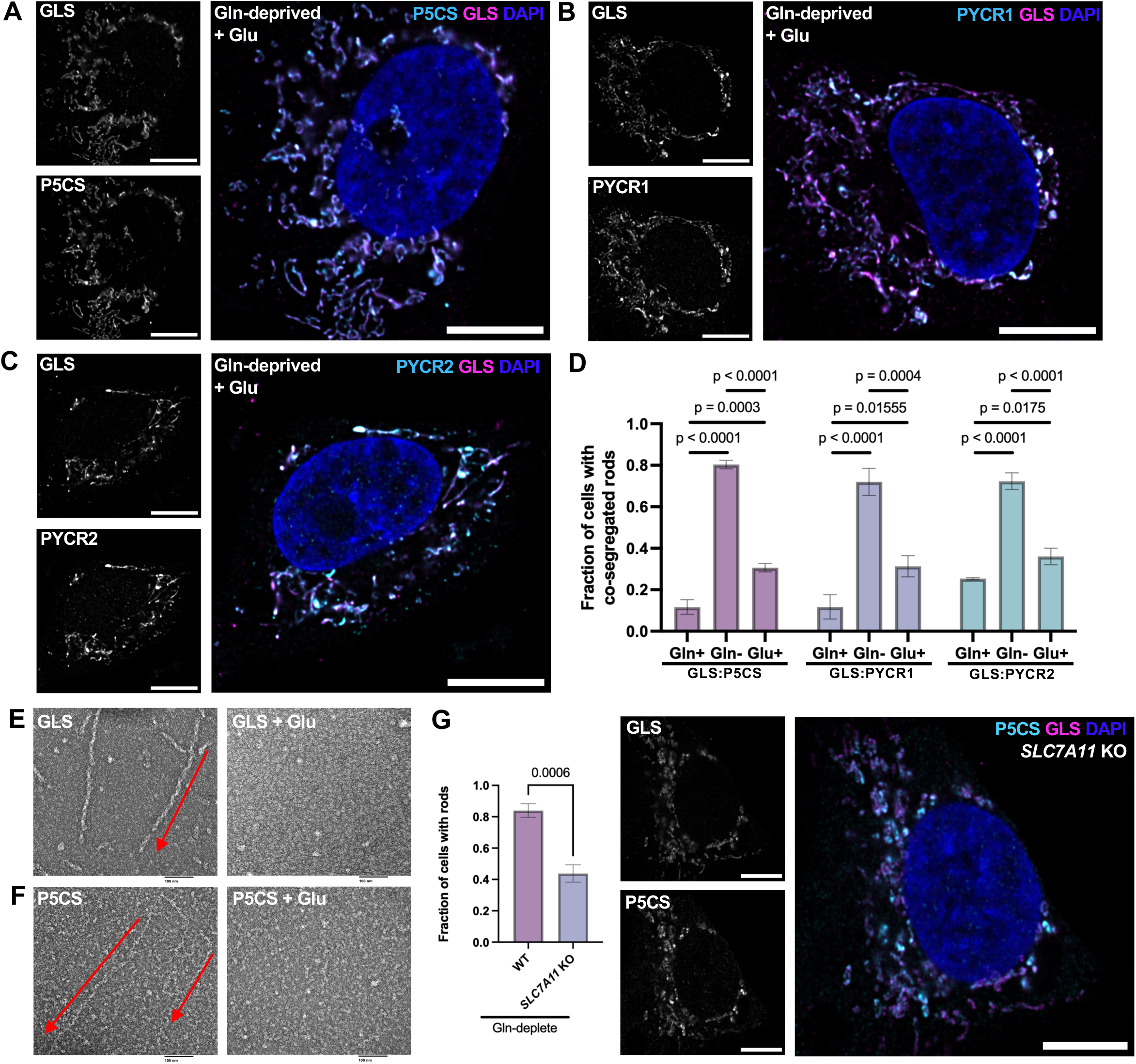
Glutamate regulates enzyme filamentation and coordinated segregation. **(A)** Supplementation with 10 mM glutamate under glutamine-deprived conditions prevents co-segregation of GLS and P5CS in Hs 578T cells. Scale bar 10 *μ*m. **(B)** Supplementation with 10 mM glutamate under glutamine-deprived conditions prevents co-segregation of GLS and PYCR1 in Hs 578T cells. Scale bar 10 *μ*m. **(C)** Supplementation with 10 mM glutamate under glutamine-deprived conditions prevents co-segregation of GLS and PYCR2 in Hs 578T cells. Scale bar 10 *μ*m. **(D)** Quantification of rod formation in glutamine-replete, glutamine-deprived, and glutamate-supplemented conditions for indicated enzyme pairs. Data from *N=*3 independent replicates. Statistical significance determined using one-way ANOVA with Tukey’s multiple comparisons test. **(E)** Purified GLS forms molecular filaments when incubated with substrate and cofactors (20 mM Gln, 50 mM inorganic phosphate), but filament formation is suppressed when excess glutamate is supplied (60 mM Glu). **(F)** Purified P5CS forms molecular filaments when incubated with substrate and cofactors (20 mM Glu, 2 mM ATP, 0.5 mM NADPH), but filament formation is suppressed at higher glutamate concentrations (60 mM Glu). **(G)** Quantification and a representative image of *SLC7A11* knockout cells under glutamine deprivation. Data from *N=*3 independent replicates. Statistical significance determined using the two-way Student’s t-test. Scale bar 10 *μ*m.

An intriguing feature of the glutamine-to-proline biosynthetic pathway is that GLS and P5CS, which catalyze consecutive reactions, are both capable of assembling into molecular filaments through the polymerization of individual enzymatic units^24–26^. We therefore investigated potential links between cellular rods and molecular filaments by testing whether glutamate availability influences GLS and P5CS filamentation *in vitro*. Negative-stain electron microscopy revealed that purified human GLS formed elongated filaments in the presence of glutamine substrate and the allosteric activator inorganic phosphate, and that these failed to form following the addition of excess glutamate (Fig. 2E). Purified P5CS also formed elongated filaments when incubated with its substrates, glutamate, ATP, and NADPH, but filaments did not form when the glutamate concentration was increased from 20 to 60 mM (Fig. 2F). Thus, corresponding with the glutamate-dependent organization of GLS and P5CS in cells, filament formation by both purified enzymes *in vitro* was sensitive to glutamate concentration. Whether the absence of P5CS filaments at higher glutamate concentrations reflects direct regulation by glutamate or an indirect consequence of catalytic turnover remains unresolved.

To investigate whether endogenous glutamate handling influences enzyme organization in cells, we next examined the role of the plasma membrane transporter SLC7A11 (xCT), which mediates glutamate export in exchange for cystine uptake and can thereby exacerbate intracellular glutamate depletion when glutamine is limiting^27–30^. We genetically deleted *SLC7A11* in MDA-MB-231 TNBC cells, which normally exhibit robust enzyme redistribution upon glutamine restriction (Supplementary Fig. 3D, 4F). Loss of SLC7A11 substantially reduced the proportion of cells containing GLS-and P5CS-positive rod-like structures during glutamine deprivation, although residual P5CS-positive puncta were again present in some cells (Fig. 2G). These findings thus support a role for endogenous glutamate availability in regulating the co-segregation of glutamine-to-proline biosynthesis enzymes during glutamine deprivation. More broadly, they support that the spatial organization of the pathway components corresponds to whether the pathway operates under nutrient-replete or nutrient-depleted conditions.

### The proline biosynthesis enzymes assemble into a metabolon associated with ATP synthase

The selective preservation of intracellular proline during glutamine deprivation, together with the co-segregation of GLS, P5CS, PYCR1, and PYCR2 into a distinct mitochondrial subpopulation, raises the possibility that these enzymes assemble into a metabolon to sustain proline biosynthesis. To determine whether these enzymes physically associate rather than simply occupy the same mitochondria, we performed co-immunoprecipitations. This revealed physical associations between GLS and each of the downstream enzymes P5CS, PYCR1 and PYCR2 (Fig. 3A). Reciprocal co-immunoprecipitations of P5CS, PYCR1 and PYCR2 further supported these associations (Fig. 3B, C). GLS did not co-precipitate with several other mitochondrial proteins, including the inner membrane protein TIMM17, matrix-localized GOT2, or the outer membrane protein TOM20 (Fig. 3D). Together, these results provide biochemical evidence that the four enzymes assemble into a multienzyme complex, which we designate a proline biosynthesis metabolon.

**Figure 3.**
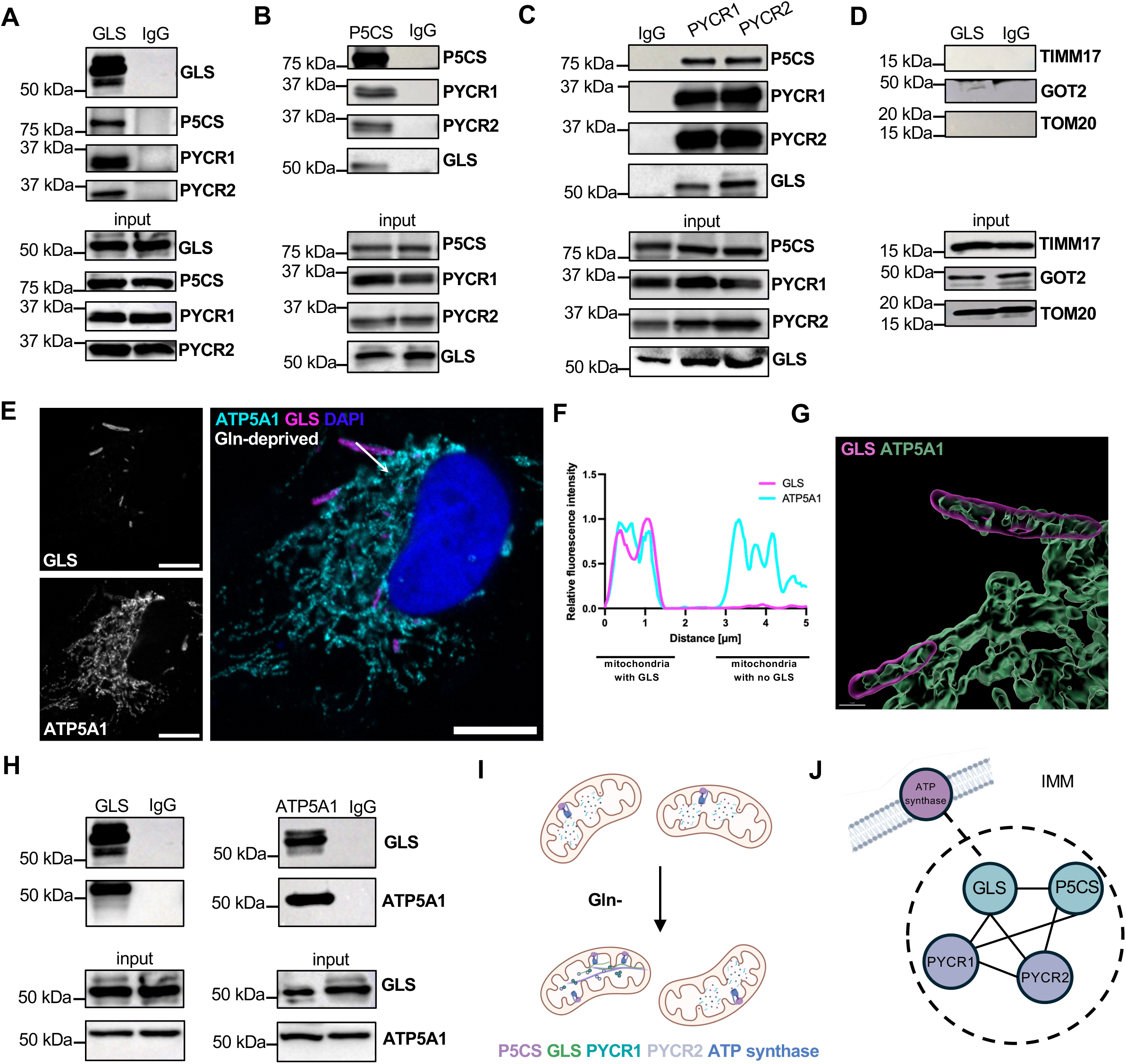
Proline biosynthesis enzymes form a metabolon that associates with ATP synthase. **(A)** Co-immunoprecipitation of GLS with partner proteins from the proline biosynthesis pathway (P5CS, PYCR1, PYCR2). **(B)** Co-immunoprecipitation of P5CS with partner proteins from the proline biosynthesis pathway (GLS, PYCR1, PYCR2). **(C)** Co-immunoprecipitation of PYCR1 and PYCR2 with partner proteins from the proline biosynthesis pathway (GLS and P5CS). **(D)** Control co-immunoprecipitations of GLS with other mitochondrial proteins: TIMM 17 (inner mitochondrial membrane marker), GOT2 (mitochondrial matrix marker), TOM 20 (outer mitochondrial membrane marker). **(E)** Representative immunofluorescence image showing the presence of ATP5A1 within GLS-enriched mitochondria under glutamine deprivation (25 *μ*M). Scale bar 10 *μ*m. **(F)** Line scan showing the spatial distribution of GLS and ATP synthase (ATP5A1) under glutamine deprivation. **(G)** 3D reconstructions showing spatial overlap between GLS and ATP synthase (ATP5A1 subunit) under glutamine deprivation. Scale bar 1 *μ*m. **(H)** Co-immunoprecipitation of GLS with ATP synthase (ATP5A1 subunit) and vice versa. **(I)** Schematic of segregation of mitochondrial subpopulations with the proline metabolon. **(J)** Schematic of the proposed interactions within the proline biosynthesis metabolon and its potential tethering to the inner mitochondrial membrane (IMM) via ATP synthase.

Previous work has identified P5CS-enriched mitochondrial subpopulations that are depleted of ATP synthase and favor the reductive biosynthesis of proline and ornithine under conditions of increased cellular reliance on oxidative phosphorylation^9^. Since our work identifies a metabolon that forms specifically when breast cancer cells experience glutamine deprivation, a condition known to restrict proliferation^31^, we probed the involvement of ATP synthase in this distinct metabolic setting. Intriguingly, immunofluorescence microscopy revealed that ATP5A1, a core subunit of ATP synthase, remained broadly distributed throughout the mitochondrial network and was readily detected within GLS-enriched mitochondria (Fig. 3E, F). Moreover, three-dimensional reconstructions showed substantial spatial overlap between GLS and ATP5A1 within the elongated mitochondrial subpopulation (Fig. 3G).

Reciprocal co-immunoprecipitations further revealed a biochemical association between GLS and ATP5A1 (Fig. 3H), supporting a physical connection between the proline biosynthesis metabolon and ATP synthase at the inner mitochondrial membrane. Based on this, we performed mitochondrial fractionation and observed that GLS and P5CS are both highly enriched in the inner mitochondrial membrane fraction (Supplementary Fig. 4G). Together, the imaging and biochemical evidence support a model in which the proline biosynthesis metabolon is tethered to the inner mitochondrial membrane via ATP synthase (Fig. 3I, J).

### The proline biosynthesis pathway supports cancer cell redox homeostasis and survival during glutamine deprivation

Having established that GLS and the downstream proline biosynthesis enzymes form a physically associated metabolon, we next investigated whether their coordinated spatial organization contributes to sustained proline synthesis during glutamine deprivation. Previous work has implicated mitochondrial dynamics, particularly DRP1-mediated fission, in the organization of metabolically distinct mitochondrial subpopulations, including those enriched in P5CS^9,16,21^. We therefore genetically depleted DRP1 and examined the organization of the metabolon enzymes during glutamine limitation. Notably, DRP1 depletion did not prevent the enzymes from extending into rod-shaped structures in parts of the mitochondrial network but markedly disrupted their co-segregation into the same mitochondrial subpopulation (Fig. 4A-4C, Supplementary Fig. 5A-5H). Three-dimensional reconstructions confirmed disrupted spatial coordination between GLS and each of the downstream enzymes P5CS, PYCR1 and PYCR2 (Fig. 4E, F). We observed matching effects following pharmacological inhibition of DRP1 (Supplementary Fig. 5I). Therefore, DRP1 depletion uncouples the concentration of individual enzymes into extended mitochondrial rods from their coordinated segregation into the same mitochondrial subpopulation.

**Figure 4.**
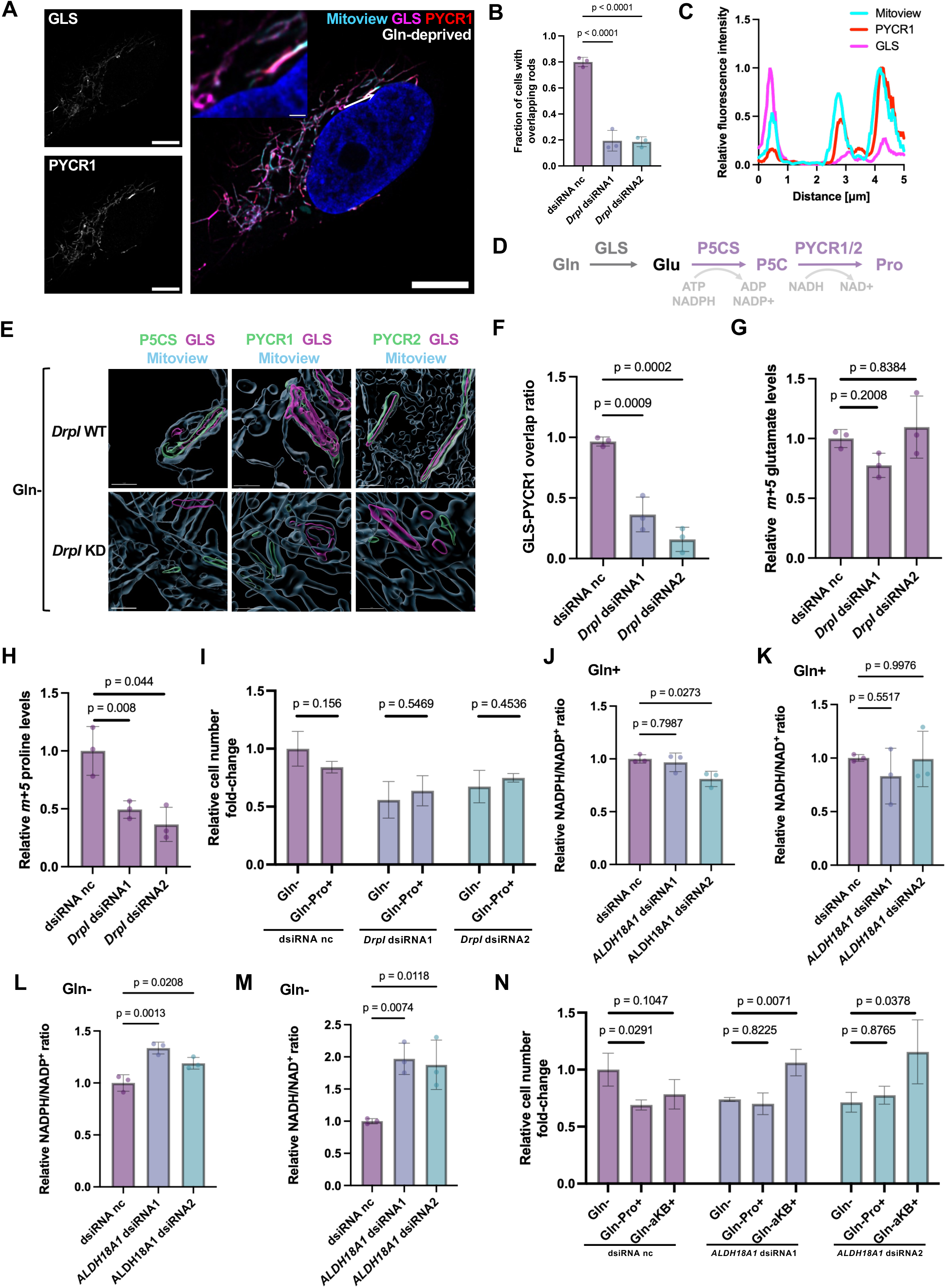
Proline biosynthesis sustains cancer cell survival during glutamine deprivation by maintaining redox homeostasis. **(A)** Representative immunofluorescence image of disrupted coordination between GLS and PYCR1 cellular rods under glutamine deprivation (25 *μ*M) in *DNM1L* (DRP1) knockdown Hs 578T cells. Scale bar 10 *μ*m. **(B)** Quantified fractions of cells with GLS-PYCR1 co-segregated rods in *DNM1L* (DRP1) knockdown Hs578T cells under glutamine deprivation. Data from *N=*3 independent replicates. Statistical significance determined using a one-way ANOVA with Tukey’s multiple comparisons test. **(C)** Line scan of PYCR1 and GLS in *DNM1L* (DRP1) knockdown Hs 578T cells. **(D)** Schematic of the proline biosynthesis pathway from glutamine, with annotated reduced cofactor consumption. **(E)** 3D reconstructions of rod coordination disruption between each pair of enzymes in *DNM1L* (DRP1) knock-down cells (lower panel) and proper coordination in wild-type cells (upper panel). Scale bar 1 *μ*m. **(F)** Quantified overlap ratio between GLS and PYCR1 signals in *DNM1L* (DRP1) knockdown cells. Data from *N=*3 independent replicates. Statistical significance determined using a one-way ANOVA with Tukey’s multiple comparisons test. **(G)** Relative levels of m+5 glutamate in *DNM1L* (DRP1) knockdown cells and scramble controls. Data normalised to internal standards, protein concentration, and then relativised to control dsiRNA group. Data from *N=*3 independent replicates statistical significance determined on log2-transformed data using a one-way ANOVA with Tukey’s multiple comparisons test. **(H)** Relative levels of m+5 proline in *DNM1L* (DRP1) knockdown cells and scramble controls. Data normalised to internal standards, protein concentration, and then relativised to control dsiRNA group. Data from *N=*3 independent replicates statistical significance determined on log2-transformed data using a one-way ANOVA with Tukey’s multiple comparisons test. **(I)** Cellular viability following 24 h exposure to glutamine deprivation or glutamine deprivation supplemented with 1 mM proline, Hs 578T cells, *DNM1L* (DRP1) knockdowns. Data normalised to levels under glutamine deprivation Data from *N=*3 independent replicates. Statistical significance determined with the two-tailed Student’s test with Welch’s correction. **(J)** Relative NADPH/NADP^+^ levels in *ALDH18A1* knockdown Hs578T cells under glutamine-replete conditions. Data normalised to negative control (dsiRNA nc). Data from *N=*3 independent replicates. Statistical significance determined with one-way ANOVA with Tukey’s multiple comparisons test. **(K)** Relative NADH/NAD^+^ ratios in *ALDH18A1* knockdown Hs578T cells under glutamine-replete conditions. Data normalised to negative control (dsiRNA nc). Data from *N=*3 independent replicates. Statistical significance determined with one-way ANOVA with Tukey’s multiple comparisons test. **(L)** Relative NADPH/NADP^+^ ratios in *ALDH18A1* knockdown Hs578T cells under glutamine-deprived conditions. Data normalised to negative control (dsiRNA nc). Data from *N=*3 independent replicates. Statistical significance determined with one-way ANOVA with Tukey’s multiple comparisons test. **(M)** Relative NADH/NAD^+^ ratios in *ALDH18A1* knockdown Hs578T cells under glutamine-deprived conditions. Data normalised to negative control (dsiRNA nc). Data from *N=*3 independent replicates. Statistical significance determined with one-way ANOVA with Tukey’s multiple comparisons test. **(N)** Cellular viability following 24 h exposure to glutamine deprivation with or without supplementation with 1 mM proline or 1 mM *α*-ketobutyrate, Hs 578T cells, *ALDH18A1* knockdowns. Data normalised to levels under glutamine deprivation. Data from *N=*3 independent replicates. Statistical significance determined with one-way ANOVA with Tukey’s multiple comparisons test.

To determine whether this loss of coordination between the metabolon components affects glutamine-derived proline synthesis, we performed stable-isotope tracing using uniformly labeled ^13^C_5_-glutamine (Fig. 4D). Following glutamine deprivation and metabolon formation, cells were exposed to a transient pulse of labeled glutamine, mimicking fluctuating nutrient availability in tumors^32^. DRP1 depletion did not significantly affect the intracellular abundance of m+5 glutamate (Fig. 4G). In contrast, the abundance of m+5 proline was substantially reduced, indicating impaired proline synthesis downstream of glutamate (Fig. 4H). Levels of m+5 glutamine were increased in DRP1-depleted cells, and no m+5 labelling of ornithine or arginine was detected (Supplementary Fig. 5J-N). These findings link disrupted coordination of the proline biosynthesis enzymes to impaired glutamine-derived proline synthesis, despite the persistence of individual enzyme-enriched mitochondrial rods.

Consistent with a role for coordinated enzyme organization in metabolic adaptation, DRP1 depletion also reduced TNBC cell viability under glutamine deprivation. However, this effect could not be rescued by supplementation with the end-product of the pathway, proline (Fig. 4I), suggesting that the importance of this pathway during glutamine deprivation might extend beyond the production of proline itself. Proline biosynthesis consumes NADPH and NADH through the sequential reactions catalyzed by P5CS and PYCR1/2, thereby regenerating oxidized NAD(P) cofactors (Fig. 4D). To determine whether this consumption of reducing equivalents contributes to cellular redox homeostasis, we depleted P5CS through knockdown of its encoding gene, *ALDH18A1* (Supplementary Fig. 5M), and measured cellular redox cofactor ratios. Under glutamine-replete conditions, P5CS depletion had little effect on either the NADPH/NADP⁺ or NADH/NAD⁺ ratio (Fig. 4J, K). In contrast, under glutamine deprivation, P5CS depletion increased both ratios, with a particularly marked effect on NADH/NAD⁺ (Fig. 4L, M).

Having established that P5CS contributes to cellular redox homeostasis during glutamine deprivation, we next investigated the importance of this function for cancer cell survival. P5CS depletion reduced TNBC cell viability under glutamine deprivation, and supplementation with proline again failed to rescue this effect (Fig. 4N). In contrast, α-ketobutyrate, an alternative electron acceptor that facilitates NAD⁺ regeneration, fully restored viability (Fig. 4N). The ability of α-ketobutyrate, but not proline, to rescue P5CS-depleted cells indicates that impaired redox homeostasis, rather than insufficient proline availability, underlies the survival defect. Collectively, our findings identify a nutrient-responsive proline biosynthesis metabolon and reveal that this pathway supports redox homeostasis and cancer cell survival when glutamine supplies are limited.

## Discussion

Enzyme filamentation has traditionally been implicated in regulating the catalytic properties of individual enzymes^13,16–18,33,34^ and, more recently, in their segregation into specialized mitochondrial subpopulations^9,35^. Our findings extend this concept to the formation of a physically associated, condition-specific metabolon comprising the filamentous enzymes GLS and P5CS^16,19,25,26,36^, along with non-filamentous PYCR1 and PYCR2^37,38^. This suggests that GLS and P5CS filamentation may contribute to their coordinated segregation with PYCR1 and PYCR2 under glutamine deprivation. We also find that proline biosynthesis is selectively sustained under these conditions. As such, filamentation may help to organize not only individual enzymes, but entire metabolic pathways into cohesive, physically associated complexes that favor specific metabolic functions when substrates become limiting.

We find that glutamate, the reaction product of GLS and the substrate of P5CS, induces disassembly of purified GLS filaments and decreases the abundance of P5CS filaments at higher concentrations *in vitro*. In cells, glutamate supplementation prevents the co-segregation of GLS, P5CS, and PYCR1/2 into a distinct mitochondrial subpopulation during glutamine deprivation. Together, these findings suggest that glutamate serves not only as a metabolic intermediate, but also as a regulatory cue linking nutrient availability to the dynamic spatial organization of the proline biosynthesis pathway.

Previous work identified a P5CS-enriched mitochondrial subpopulation that is depleted of ATP synthase under conditions of increased reliance on oxidative phosphorylation^9^. In contrast, the GLS-enriched mitochondrial subpopulation that forms during glutamine deprivation retains ATP synthase, whose core subunit ATP5A1 co-immunoprecipitates with GLS. This biochemical association raises the possibility that ATP synthase helps anchor the proline biosynthesis metabolon to the inner mitochondrial membrane. Such an arrangement could also position P5CS near a local source of ATP, which is required for its catalytic activity^36,39^. These observations suggest that the spatial relationship between mitochondrial enzyme filamentation and ATP synthase can vary across metabolic contexts.

Beyond these molecular associations, mitochondrial dynamics also contribute to the reorganization of mitochondrial metabolism^12,35,40^. We show that depletion of the mitochondrial fission regulator DRP1 does not prevent individual pathway enzymes from concentrating into elongated mitochondrial rods but markedly disrupts their coordinated segregation into the same mitochondrial subpopulation. This loss of coordination is accompanied by reduced incorporation of glutamine-derived carbon into proline, without a significant change in the abundance of labeled glutamate. Although DRP1 depletion may have broader consequences for mitochondrial physiology, these findings support a functional role for coordinated enzyme segregation in sustaining proline biosynthesis during glutamine deprivation.

Proline biosynthesis has previously been implicated in the dissipation of excess mitochondrial reducing equivalents under conditions of elevated metabolic demand^41,42^. The successive reactions catalyzed by P5CS and PYCR1/2 consume NADPH and NADH, respectively^43,44^, linking proline synthesis to the regeneration of oxidized redox cofactors. Here, we find that this function becomes particularly important during glutamine deprivation, when P5CS depletion markedly increases both NADPH/NADP⁺ and NADH/NAD⁺ ratios, despite having little effect under glutamine-replete conditions. Importantly, α-ketobutyrate, an alternative electron acceptor that facilitates NAD⁺ regeneration^45^, restored the viability of P5CS-depleted cells under glutamine deprivation, whereas proline supplementation did not. These findings indicate that the contribution of proline biosynthesis to cancer cell survival under these conditions reflects its role in maintaining redox homeostasis rather than simply supplying proline. This contrasts with a previous report that P5CS suppression can promote the proliferation of stomach and prostate cancer cells by conserving glutamate for *de novo* glutamine synthesis^46^, highlighting the context-dependent importance of proline biosynthesis for cancer cell fitness. More broadly, our findings illustrate how the physiological significance of a biosynthetic pathway can extend beyond its end product to the biochemical reactions involved in its synthesis. They further implicate nutrient-responsive enzyme filamentation in the coordinated spatial organization of entire metabolic pathways, extending its potential role beyond the regulation of individual enzymes to the preservation of selected metabolic functions under nutrient scarcity.

## Acknowledgments

We thank Dr. Leemor Joshua-Tor, Dr. Leah Braviner and Stephanie Goldsmith for supplying the Sf9 insect cell line and relevant reagents for protein expression, Dr. Ankit Tiwari for supplying the mitoBFP2-expressing construct, and Dr. Linda Van Aelst, Dr. Nicholas Tonks, Dr. Tobias Janowitz, and Dr. Gabrielle Pouchelon for discussions. We thank the Microscopy Shared Resource and the St. Giles Advanced Microscopy Center, the Flow Cytometry Shared Resource and the Mass Spectrometry Shared Resource at Cold Spring Harbor Laboratory, which are supported by the Cancer Center Support Grant 5P30CA045508 from the National Institutes of Health. We also thank the Proteomics and Metabolomics Core Facility at H. Lee Moffitt Comprehensive Cancer Center & Research Institute, Tampa, Florida; supported by the grant P30-CA076292 from National Institutes of Health. Purchase of the LSM980 Airyscan 2 was supported by the grant 1S10OD034372-01 from the National Institutes of Health.

## Author Contributions

Conceptualization: NK, MJL

Methodology: NK, SY, SP, T-LW, W-HY, PC, MJL

Investigation: NK, SK, JDRS, YQ, SJ, SP, PC, T-LW

Visualization: NK, T-LW, MJL

Funding acquisition: NK, MJL Supervision: MJL

Writing – original draft: NK, SK, JDRS, MJL

## Data, code, and materials availability

All relevant data are available in the main text and in the supplementary materials. Relevant reagents will be made available by request to the corresponding author.

## Supplementary Material

### Materials and Methods

#### Cell culture

All breast cancer cell lines (MDA-MB-231, Hs 578T, BT-20, HCC 1806, JIMT-1, MDA-MB-231 BrM) were maintained in Dulbecco’s Modified Eagle Medium (DMEM, Gibco) or Roswell Park Memorial Institute Medium (RPMI, for lentiviral transductions) supplemented with 10% fetal bovine serum (FBS). Experimental media was made in glutamine-free DMEM (Gibco), supplementing with appropriate amino acids in indicated concentrations (proline, dimethyl oxoglutarate, dimethyl 2-oxobutyrate, aspartate, glutamate, glutamine, all from Sigma-Aldrich). Cells were routinely verified to be mycoplasma-free using the MycoAlert Detection Kit (Lonza).

For proliferation and viability assays, cells were seeded in 6-well plates at 50 000 cells/well or 100 000 cells/well and in triplicates per condition. After overnight attachment, cells were washed briefly in phosphate buffer saline (PBS) and exposed to experimental media as indicated for individual experiments. For normalization, a control plate was counted at the moment of exposure to experimental media. For cell counting, cells were first trypsinized (0.05% dissociation reagent-EDTA, Gibco) and stained with trypan blue (Corning) for viable count determination (CellCount). Alternatively, cells were seeded at 2500 cells/well in a 96-well plate, allowed to attach overnight and then incubated in experimental medium as indicated for individual experiments. Viability was read using CCK-8 cell viability assay kit (Vazyme). For inhibition experiments, DRP1i27 (MedChemExpress) was used at a final concentration of 10*μ*M, and added to experimental media for plated cells in ibidi plates for immunofluorescence analysis as described below.

#### Antibodies

Unless otherwise indicated, all antibodies were used at a 1:1000 dilution for western blot, at 1:400 for immunofluorescence, and at 5*μ*g per co-immunoprecipitation reaction. The following antibodies were used in this study: anti-GLS1 (Cell Signaling Technology, 56750, for western blot co-immunoprecipitation, and at 1:500 for immunofluorescence), anti-KGA/GAC (Proteintech, 66265-1-IgG, 1:250 for immunofluorescence, for western blot), anti-P5CS (Proteintech, 17716-1-IgG, for western blot, co-immunoprecipitation, and immunofluorescence), anti-P5CS (Proteintech, 68184-1-Ig, 1:300 for immunofluorescence), anti-PYCR1 (Proteintech, 13108-1-AP, for western blot, co-immunoprecipitation, and immunofluorescence), anti-PYCR1 (Proteintech, 66510-1-Ig, for western blot and immunofluorescence), anti-PYCR2 (Proteintech, 68488-1-Ig, for western blot and immunofluorescence), anti-PYCR2 (Proteintech, 17146-1-AP, for western blot, coimmunoprecipitation, and immunofluorescence), anti-Hsp60 (Proteintech, 15282-1-AP, for immunofluorescence), anti-ATP5A1 (Proteintech, 66037-1-IgG, for western blot, co-immunoprecipitation, and immunofluorescence), anti-ATP5A1 (Proteintech, 14676-1-AP, for western blot), anti-GOT2 (Abcam, ab171739, western blot, immunofluorescence), anti-ACO2 (Cell Signaling Technology, 6571, 1:400 for immunofluorescence, for western blot), anti-TIMM17a (Cell Signaling Technology, 4594, for western blot), anti-TOM20 (Cell Signaling Technology, 42406, for immunofluorescence and western blot), anti-HSPE1 (AB Clonal, A5580, for western blot), anti-SLC7A11 (Cell Signaling Technology, 12691, for western blot), anti-TOMM40 (Proteintech, 18409-1-AP, for western blot), anti-DRP1 (Cell Signaling Technology, 8570, for western blot) normal rabbit IgG (Cell Signaling Technology, 2729, for co-immunoprecipitation), normal mouse IgG (Cell Signaling Technology, 68860, for co-immunoprecipitation).

##### Lentiviral transduction

Lentivirus transduction was used to generate knockouts of endogenous *GLS*, knockouts of endogenous *SLC7A11*, and stably expressing cell lines of GFP-tagged GAC, the most abundant splice variant of GLS in cultured breast cancer cells^5^, and mitochondrial BFP2. Plasmid expressing mBFP2 was a kind gift of Dr. Ankit Tiwari. In brief, lentiviral particles were generated in HEK-293T cells by combining 500 *μ*l Opti-MEM (gibco) with 5 *μ*g of the appropriate DNA construct, 5 μg pCMVΔR8.2, and 1 μg pMD2.G; and combining 500 *μ*l Opti-MEM (Corning) with 32 *μ*l of 1 mg/mL PEI 25K (Polysciences). Aliquots were incubated separately at room temperature for 5 minutes and then combined for 15 minutes before dropwise addition to HEK-293T cells grown in 10 cm dishes. Virus-containing supernatant was collected at 48 h and 72 h, combined, and filtered with a 0.45 *μ*m filter. Virus-containing supernatant was diluted 1:3 in fresh cell culture medium containing 10 *μ*g/mL polybrene, applied to cells, and incubated for 8 hr at 37°C and 5% CO_2_. The medium was then substituted with fresh RPMI supplemented with 10% diFBS (Cytiva). Cells were incubated for additional 48 hr before addition of antibiotic selection (1*μ*g/mL puromycin dihydrochloride (VWR) for knockouts or 500 *μ*g/mL hygromycin B (VWR) for expression constructs). Selection was applied for 10 to 14 days with passaging, after which single clones were selected by single-cell plating in 96-well plates and cultured with passaging until validation of knockouts by targeted sequencing of the genomic locus and western blot.

Guide RNAs for *GLS* knockouts were cloned into pLentiCRISPRv2. Sequences of guide RNAs are as follows:

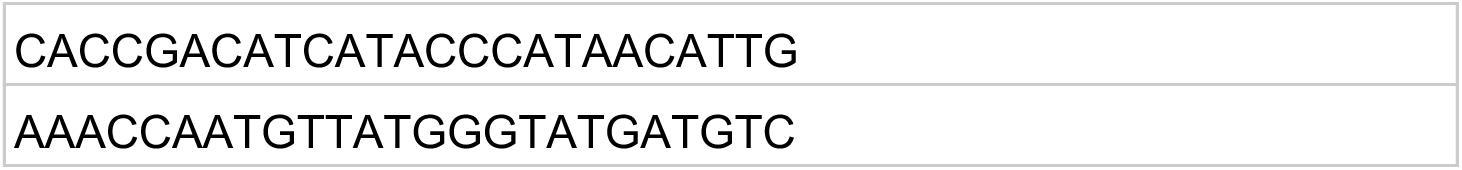

Guide RNAs for *SLC7A11* knockouts were cloned into pTLCv2, an inducible KO plasmid based on pLentiCRISPRv2.

Sequences of guide RNAs are as follows:

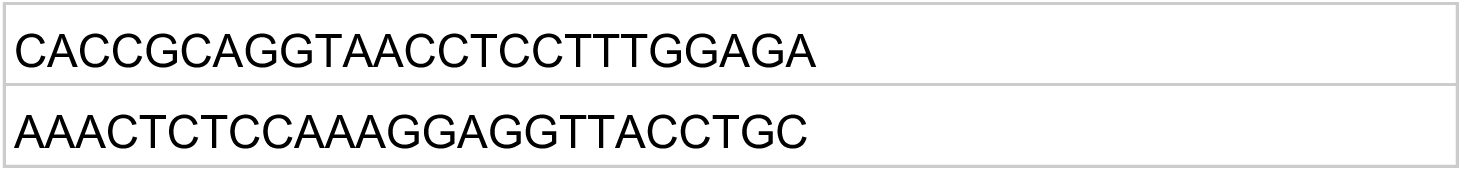

##### Immunofluorescence

Cells were seeded at 2-4×10^4^cells/well in 8-well Ibidi microslides ibidiTreat (Ibidi) and after overnight attachment treated with experimental medium as indicated for individual experiments. For visualising mitochondria, cells were treated with 100nM-200nM Mitoview 640 Fix (Biotium) for three hours. Samples were then fixed in 4% paraformaldehyde (Electron Microscopy Sciences) for 10 minutes or in 90% methanol (VWR) for 66037-1-IgG antibody. Samples were permeabilized with 0.1% Triton X-100 in PBS for 10 minutes and blocked in 3% FBS in PBS-0.05% Tween-20 for 1 hour. Antibodies were incubated at indicated dilutions overnight at 4°C. After incubation, samples were washed three times for 10 minutes in PBS-0.05% Tween-20 and incubated in secondary antibodies (goat anti-mouse Alexa Fluor 488, goat anti-rabbit Alexa Fluor 488, goat anti-mouse Alexa Fluor 647, goat anti-rabbit Alexa Fluor 647, goat anti-rabbit Alexa Fluor 568, all Invitrogen) and 1:1000 dilution of 1mg/ml DAPI (Invitrogen). Finally, samples were washed three times for 10 minutes in PBS-0.05% Tween-20 and left in PBS for image acquisition on the Carl Zeiss LSM 980+Airyscan 2 as described below.

##### Immunofluorescence on tumor samples

Breast cancer tumour samples (ID: FR0001CECE) were purchased from OriGene Technologies, cut using a Leica CM3050s cryostat into 10 μm sections and mounted on Superfrost slides. Immunofluorescence protocol from above was adapted for staining of the on-slide specimens using the same reagents and buffer composition. Samples were mounted using SlowFade Glass Soft-Set AntiFade Mountant (Thermo Fisher Scientific) and 1.5 coverslips and imaged on the Carl Zeiss LSM 980+Airyscan 2 as described below.

##### Image acquisition and analysis

Microscopy data were acquired in the Airyscan2 Multiplex or Super Resolution mode using an inverted LSM 980 + Airyscan2 confocal laser scanning microscope with Plan-Apochromat 20x/0.8NA air or Plan-Apochromat 63x/1.4NA oil immersion objective (Carl Zeiss Microscopy, White Planes, NY). DAPI was excited with a 405 nm solid-state diode laser, and antibody-conjugated fluorophores were excited using 488 nm, 561 nm, or 639 nm solid-state diode lasers, as appropriate. Sequential frame scanning and appropriate filter settings were used to minimize crosstalk between fluorescence channels. Images of individual filaments in mitochondria were acquired in the Airyscan super-resolution mode at additional 12x zoom and eightfold averaging. Fluorescence signals were detected using 32-channal array GaAsP-PMT Airyscan detector. All data were processed in the Zeiss Zen Blue 3.12 software (Carl Zeiss Microscopy, White Planes, NY) using the 3D Airyscan Joint Deconvolution Processing with quality threshold set to 0 and 10 iterations.

Surface function was used for volumetric reconstruction of signals using the absolute intensity thresholding was applied to all channels, and the thresholding cut-off was optimized manually based on the fluorescence intensity of each fluorescence channel. Animations for supplementary material were prepared in Imaris using the Animation function combining the representative frames at 10 frames per second frame rate. Line intensity profiles were obtained in Zen 3.12 on unprocessed images. Quantification of filament abundance was done in Volocity 7 (Quorum Technologies, ON, Canada) by reconstruction of nuclear volumes from DAPI signal and applying automated intensity thresholding and size parameters of at least 0.5*μ*m for objects to identify rods of either protein. Rods were assigned to nuclei based on proximity and cells were binned into rod-containing or pan-mitochondrial categories based on whether any objects were identified. For filamentous rod coordination, rod signal for every protein of interest was reconstructed into area in Imaris 11.0.1 using the Surface Reconstruction function on 20x aerial images featuring an appropriate number of cells (larger than 100 for each quantification). Cells were binned into coordinated or non-coordinated based on area overlap of signals for the protein pair being larger than 70% for coordination. Data plotting and statistical testing were conducted using Prism 9. For representative purposes in figures, appropriate sections from 3D stacks were prepared in Fiji. No quantification was performed on post-processed images, and fluorescence intensity was not used as a proxy for protein expression levels. To ensure consistency in intensity measurements within and between datasets, standardized imaging and quantitative analysis parameters were applied throughout the project.

##### Western blot

Cells were plated in 6-cm dishes and harvested at 70% confluency with RIPA buffer (Sigma-Aldrich) supplemented with Pierce MiniProtease Inhibitor Tablets (Thermo Fisher Scientific). Input was normalized through protein concentration detected in Pierce BCA protein assay (Thermo Fisher Scientific). Samples were loaded on Novex Tris-Glycine Mini Protein Gels (Thermo Fisher scientific) and transferred onto a 0.45*μ*m nitrocellulose membrane (Thermo Fisher Scientific) using Mini Blot modules (Invitrogen). Membranes were blocked in 5% non-fat milk for an hour at room temperature and incubated in primary antibodies overnight at 4°C as described above. Membranes were washed in tris buffered saline-0.05% Tween-20 three times for 10 minutes, incubated with secondary antibodies for 30 minutes at room temperature (anti-mouse HRP-conjugated secondary antibody and anti-rabbit HRP-conjugated secondary antibody, Cell Signaling Technology), then washed again and imaged using SuperSignal West Pico Plus chemiluminescent substrate (Thermo Fisher Scientific) on Azure 300 gel imager.

##### Co-immunoprecipitation

Cells were grown in 15-cm dishes to 80% confluency and incubated in experimental medium for 24 hr as indicated for individual experiments. Before harvesting, cells were treated with 2mM dithiobis(succinimidylpropionate) (DSP, Lomant’s reagent, Thermo Fisher Scientific) to crosslink filaments for 30 minutes as per manufacturer’s instructions. Lysates were then obtained using RIPA buffer (Sigma-Aldrich) supplemented with Pierce MiniProtease Inhibitor Tablets (Thermo Fisher Scientific). Antibodies as indicated above were incubated for 1 hr at room temperature in the same RIPA buffer and Dynabeads G (Thermo Fisher Scientific) as per manufacturer’s instructions. Input was normalized through protein concentration detected in Pierce BCA protein assay (Thermo Fisher Scientific). Normalized lysates were incubated with primary antibodies overnight at 4°C (for proteins of interest, and IgG control antibodies for their respective species, and non-antibody conjugated beads as negative controls). Beads were washed two times using a high-salt containing buffer (50 mM Tris-HCl, pH 7.4; 1 M sodium chloride, 1 mM EDTA, 1% NP-40 substitute, 0.5 % sodium deoxycholate) and twice using a low-salt containing buffer (20 mM Tris-HCl pH 7.4, 10 mM magnesium chloride) on a magnetic rack. Co-precipitated proteins were eluted in 1x Laemmli buffer in low-salt containing buffer for 15 minutes by boiling at 95°C, DSP was then cleaved with 50mM DTT at 37°C for 30 minutes. Fractions and inputs were analyzed by western blot as described above.

##### Cellular fractionation and mitochondrial isolation

Cells were grown in 15-cm dishes to 80% confluency and harvested by trypsinization, washed in PBS and then resuspended in 1.1mL of ice-cold RSB hypotonic buffer (10 mM NaCl, 1.5 mM MgCl_2_, 10 mM Tris-HCl – pH=7.5) and allowed to swell for 10 minutes. Cells were mechanically lysed with a Dounce homogenizer and supplemented with 0.8mL 2.5× MS homogenization buffer (525 mM mannitol, 175 mM sucrose, 12.5 mM Tris-HCl pH=7.5, 2.5 mM EDTA pH=7.5). Unbroken cells and nuclei were removed at 1300 g for 5 minutes. Supernatants were centrifuged at 10000 g for 15 minutes to separate mitochondrial fraction (pellet) from cytoplasm (supernatant). Pellet was resuspended in 1x MS buffer (210 mM mannitol, 70 mM sucrose, 5 mM Tris-HCl pH=7.5, 1 mM EDTA pH=7.5) and subjected to fluorescence-activated sorting or mitochondrial sub-fractionation as described below.

##### Mitochondrial sub-fractionation

Mitochondrial sub-fractionation was adapted from previously described work (*47*). In brief, isolated mitochondria were resuspended in 100 mM sodium carbonate and 2 mM phenylmethylsulfonyl fluoride for 30 minutes on ice to lyse mitochondria and strip the membrane of any membrane-associated, but not membrane-integral proteins. Unlysed mitochondria were pelleted at 15 000g for 2 minutes at 4°C, and the supernatant was ultracentrifuged for 40 minutes at 75 000g at 4°C. Supernatant was briefly washed with 1x wash buffer ((50 mM Tris-HCl pH=7.4, 275 mM sodium chloride, 20 mM potassium chloride, 14 mM disodium hydrogenphosphate) and centrifuged at 20 000g for 5 minutes at 4°C three times. Resuspended pellet contains membrane-integral proteins, whereas the supernatant contains all soluble and membrane-associated proteins. Another fraction of mitochondria was assayed by sonication, which disrupts the mitochondrial membranes without stripping membrane-associated proteins. Following sonication, unlysed mitochondria were pelleted at 15 000 g for 2 minutes at 4°C, and the supernatant was ultracentrifuged for 40 minutes at 75 000g at 4°C. Supernatant was briefly washed with 1x wash buffer and centrifuged at 20 000g for 5 minutes at 4°C three times. Resuspended pellet contains membrane-integral and membrane-associated proteins, while the supernatant contains soluble non-membrane-associated mitochondrial proteins.

Another fraction of mitochondria was assayed by proteinase K (New England Biolabs) treatment and Triton X-100 (Sigma Aldrich). First, undisrupted isolated mitochondria were assayed with 10 U/mL proteinase K, which digests any external proteins. Second, undisrupted isolated mitochondria were assayed with 5% Triton X-100 to lyse the mitochondria, which releases all mitochondrial proteins. Third, undisrupted isolated mitochondria were assayed with 10 U/mL proteinase K and 5% Triton X-100, which digests all mitochondrial proteins. Fourth sample was supplemented with the appropriate amount of water as a control. Samples were stored at room temperature for 30 minutes and supplemented to 2 mM phenylmethylsulfonyl fluoride. All fractions were analyzed to probe presence of proline metabolon filament-forming proteins using western blot as described above.

##### Fluorescence-activated mitochondrial sorting assay

Isolated mitochondria in suspension were sorted using a BD FACSymphony S6 cell sorter equipped with a 100 µm nozzle at a sheath pressure of 20 psi. Instrument control and sorting were performed using BD FACSDiva software (version 9.9), with the 3+ Way Purity sort mode. Before acquiring mitochondrial samples, 2.04 µm and 5.05 µm reference beads (Spherotech, SPHERO; catalog nos. PPS-1 and PPS-3, respectively) were run to guide the approximate placement of the light-scatter gate.

Fluorescence compensation between mBFP2 and GFP was calculated automatically in BD FACSDiva using GFP-only, BFP-only, and unstained mitochondrial controls. Fluorescence gates were established using the unstained control to define background fluorescence and the single-positive controls to define positive and negative populations. Four-way sorting was performed to collect BFP⁺GFP⁺, BFP⁺GFP⁻, BFP⁻GFP⁻, and BFP⁻GFP⁺ fractions into collection tubes containing 4% sodium dodecyl sulfate (SDS) in 1x MS buffer, and snap-frozen in liquid nitrogen before western blot analysis.

##### Metabolomics sample preparation and data acquisition

Cells were seeded in 6-well plates, grown to 80% confluency, and pre-treated in experimental medium as indicated for individual experiments. Tracing with ^13^C_5_-glutamine and ^13^C_6_-glutamine (Cambridge Isotope Laboratories) was applied as indicated for individual experiments. For pulse experiments, cells were treated for 23 hours in glutamine-deprived DMEM (gibco) supplemented to 25μM glutamine, and then exposed to glutamine-deprived DMEM supplemented with 1 mM tracer for 1 hr. After tracer exposition, the medium was quickly aspirated, and metabolism was quenched by adding 500 μL of extraction solvent consisting of 4:4:2 ACN:H_2_O:MeOH supplemented with 85 μM metabolomics N- and C-labelled amino acid mix as internal standards (MSK-A2-1.2, Cambridge Isotope Laboratories). Cells were harvested into the extraction solvent and incubated on dry ice for 20 minutes, after which cell debris was pelleted at 16 000 g for 10 minutes. Supernatants were dried down in a vacuum centrifuge and then resuspended immediately before LC-MS analysis in 100 μL of 5:3:2 ACN:H_2_O:MeOH. Samples were vortexed, sonicated for 5 minutes, and centrifuged at 17,000 × g and 4 °C for 3 minutes before being transferred to autosampler vials for LC-MS analysis.

LC-MS runs were conducted at the Cold Spring Harbor Laboratory Mass Spectrometry Shared Resource and at Proteomics and Metabolomics Core Facility at H. Lee Moffitt Comprehensive Cancer Center & Research Institute, Tampa, Florida. The mass spectrometer was preliminarily calibrated in both positive and negative polarity with the Pierce LTQ Velos ESI Positive Ion Calibration Solution (PN 88323) and the Pierce ESI Negative Ion Calibration Solution (PN 88324) respectively. Prior to sample analysis, a system suitability test (SST) was conducted on the QExactive HF mass spectrometer (Thermo Fisher Scientific) to evaluate chromatographic performance. Metabolites were assessed for peak shape, resolution, and mass error stability over time, and isobaric amino acids leucine and isoleucine were checked for baseline chromatographic peak resolution (confirmed), and mass error was < 5 ppm. For polar metabolite analysis with hydrophilic interaction liquid chromatography (HILIC), the system suitability test was conducted analyzing a standard cell extract (HEK 293 cells).

LC-MS analysis was performed using a Vanquish Horizon UHPLC coupled to a QExactive HF mass spectrometer (Thermo Fisher Scientific), operating in heated electrospray (HESI) ionization. HILIC chromatographic separation was completed with a XBridge BEH Amide column (2.1 x 100 mm, 2.5 µm, Waters) kept at 40 °C. Mobile phase A consisted of 20 mM NH_4_-acetate in 90:10 H2O:ACN with 0.05% medronic acid. Mobile phase B consisted of 2 mM NH_4_-acetate in 90:10 ACN:H2O with 0.05% medronic acid. After loading each sample, analytes were resolved on a 25 min step gradient as follows: 0-12.5 min 100% → 90%B; 12.5-19 min 90% → 60%B; 19-20 min 60%B; 20-25 min 100%B. Sample injection was randomized to reduce batch effects, with the injection volume set to 5 µL. Profile data was collected from 70-700 m/z in polarity switching mode (+3.5 kV to -2.5 kV) at 120,000 resolution and with AGC target set to 1E6 and a maximum IT of 50 ms. The capillary temperature was set to 300 °C with a sheath gas rate of 50, aux gas of 10, and a sweep gas of 0. Spectra were analyzed in Skyline 26 against retention times obtained in a spectral library under same acquisition conditions with gating applied across all isotopologues. For assessing statistical significance, data were analyzed in Prism (add version), log2-transformed and interrogated by using the unpaired Student’s t-test with Welch’s correction.

##### Protein expression and purification

Truncated Δ71-GAC was expressed in BL-21 cells (New England Biolabs) using a 6-His-tagged pET28a vector. In brief, construct was transformed into BL-21 cells, which were grown on Luria-Bertani agar and 50 *μ*g/mL kanamycin overnight. A single colony was then picked and propagated overnight in 10 mL of LB broth and kanamycin and expanded to 1L of expression culture. Cells were induced with 0.5 mM IPTG at OD=0.5 overnight at 25°C and spun down the following morning. Pellets were lysed in 100 mM Tris-HCl, pH=8, 300 mM NaCl, 0.5mM Tris(2-carboxyethyl)phosphine (TCEP), and supplemented with Pierce MiniProtease Inhibitor tablets (Thermo Fisher Scientific), lysozyme, DNase I, and glycerol for final concentration of 5%. Cells were lysed by sonication, after which unlysed cells were removed in a brief low-speed spin. Membranes were separated from the cytosolic fraction by ultracentrifugation for 45 minutes at 40 000 g using the SW-28 swinging bucket rotor (Beckman Coulter). Protein was first purified through affinity chromatography with a 1 hr incubation at 4°C with Ni-NTA resin (Cytiva), and then further with anion exchange chromatograph using the HiTrap Q HP column (Cytiva) on an automated AKTA Purifier purification system. Protein was concentrated using a 30 kDa concentrator (Sigma Aldrich) and frozen at -80°C until further use.

Human full-length P5CS with a C-terminal strep tag was expressed using the baculovirus expression system adapted from previous work (*48*). In brief, baculovirus particles containing the full-length gene was generated using engineered DH-10Bac cells that were a kind gift of Drs. Leah Braviner and Leemor Joshua-Tor. Baculovirus was isolated from positive controls as described before and virus was generated using Sf9 insect cells maintained in HyClone CCM3 cell culture media (Cytiva) at 28°C. Cells were first transfected in adherent culture at 6-well plates using XtremeGENE 360 (Roche) as the transfection reagent and then scaled up to third generation of the virus. Third viral passage was used to infect Sf9 cells in culture at 2.5×10^6^ cells/mL at 28°C for 48 hours, after which cells were spun down and subject to purification. In brief, pellets were resuspended in a lysis buffer containing 200 mM NaCl, 25 mM Tris/HCl, pH=8, and 0.5 mM TCEP, supplemented with DNase I, 1mM phenylmethylsulfonyl fluoride, and Pierce MiniProtease Inhibitor tablets. Cells were lysed by sonication, spun down to remove unbroken cells, and subject to ultracentrifugation for 45 minutes at 40 000 g using the SW-28 swinging bucket rotor (Beckman Coulter). Lysates were incubated with Strep resin (IBA Life Sciences) for 1 hr at 4°C, washed extensively, and eluted with 25mM Tris-HCl pH=8, 200 mM NaCl, 0.5 mM TCEP, and 7mM desthiobiotin. Elutions were concentrated and further purified by size-exclusion chromatography using a Superdex 200, 10/300 gel filtration column (Cytiva) and AKTA Purifier automated purification system.

##### Transmission Electron Microscopy

Purified protein at final concentration of 1 *μ*M was incubated in 25 mM Tris/pH=7.4, 200 mM NaCl and 0.5 mM TCEP and supplemented with amino acids as indicated for individual experiments. Samples were incubated for 1 minute on previously glow-discharged carbon film 300 mesh grids (Electron Microscopy Sciences). Excess sample was blotted off, and samples were then washed for 30 seconds by lightly placing the grid against a drop of 25 mM Tris/pH=7.4. Sample was then stained for 1 minute on uranyl acetate using the same method as for the wash. Excess stain was blotted off, and the sample was allowed to air-dry. Samples were imaged at 100 kV using a Hitachi HT7800 transmission electron microscope equipped with an AMT MkII lens 15-megapixel digital camera.

##### Knockdowns

Transient knockdowns were generated in Hs 578T cells by using custom dsiRNA for genes of interest (Integrated DNA Technologies). Sequences of dsiRNA for *ALDH18A1* are: rGrUrC rArCrA rArArA rArUrC rGrArU rGrGrA rArUrC rArUT T (dsiRNA1, *ALDH18A1* 13.1) and rCrUrGrArArGrUrGrG rGrArArUrCrArGrUrArCrArUrCrGAG (dsiRNA2, *ALDH18A1* 13.2). For DRP1, rCrArU rGrUrU rUrCrA rCrArA rGrArA rGrArU rArArA rCrGG A (dsiRNA1, *DRP1* 13.1) and rGrArU rUrCrA rUrArC rUrUrA rArArC rCrUrG rArArA rGrCA G (dsiRNA2, *DRP1* 13.2) were used. Non-targeting dsiRNA (RNA nc) was used as a control (51-01-14-04, Integrated DNA Technologies).

In brief, 9 *μ*L lipofectamine RNAiMax (Thermo Fisher Scientific) was mixed with 150 *μ*L opti-MEM (Gibco); and 3 *μ*L of 10 *μ*M dsiRNA in RNA duplex buffer (Integrated DNA technologies) was mixed with 150 *μ*L opti-MEM. Mixtures were pooled and incubated for 5 minutes at room temperature, after which they were added dropwise to cells in fresh culture medium in 6-well plates. Efficiency of knockdowns was verified by western blot, and effect on mitochondrial morphology for the *DNM1L* knockdown was verified by immunofluorescence as described above.

##### NAD(P)H/NAD(P) ratio measurements

Levels of NADH and NAD^+^ were measured using the colorimetric kit from MedChemExpress according to manufacturer’s instructions. Levels of NADPH and NADP^+^ were measured using the colorimetric kit from the same manufacturer. In brief, cells were grown to 80% confluency in six-well plates, and metabolites extracted using the manufacturer’s extraction solvent (200 *μ*L) by gentle scraping. Lysates were incubated for 10 minutes on ice, and the non-soluble fraction was pelleted at 12000 g for 10 minutes at 4°C and filtered through a 10 kDa protein concentrator. For extraction of the reduced forms, half of the extracted sample was heated at 60°C for 30 minutes. Samples were incubated with the reaction reagent provided by the manufacturer containing an appropriate enzyme for readout of either metabolite. Spectroscopic values were first corrected for no-sample background controls, and then levels of NADH were subtracted from total NAD^+^ and NADH to obtain levels of NAD^+^, and the ratio was obtained by dividing the two. Values were then normalized against the ratios obtained for the control cell lines. Statistical analyses were conducted in Prism 9.

---

**Supplementary Figure 1.**
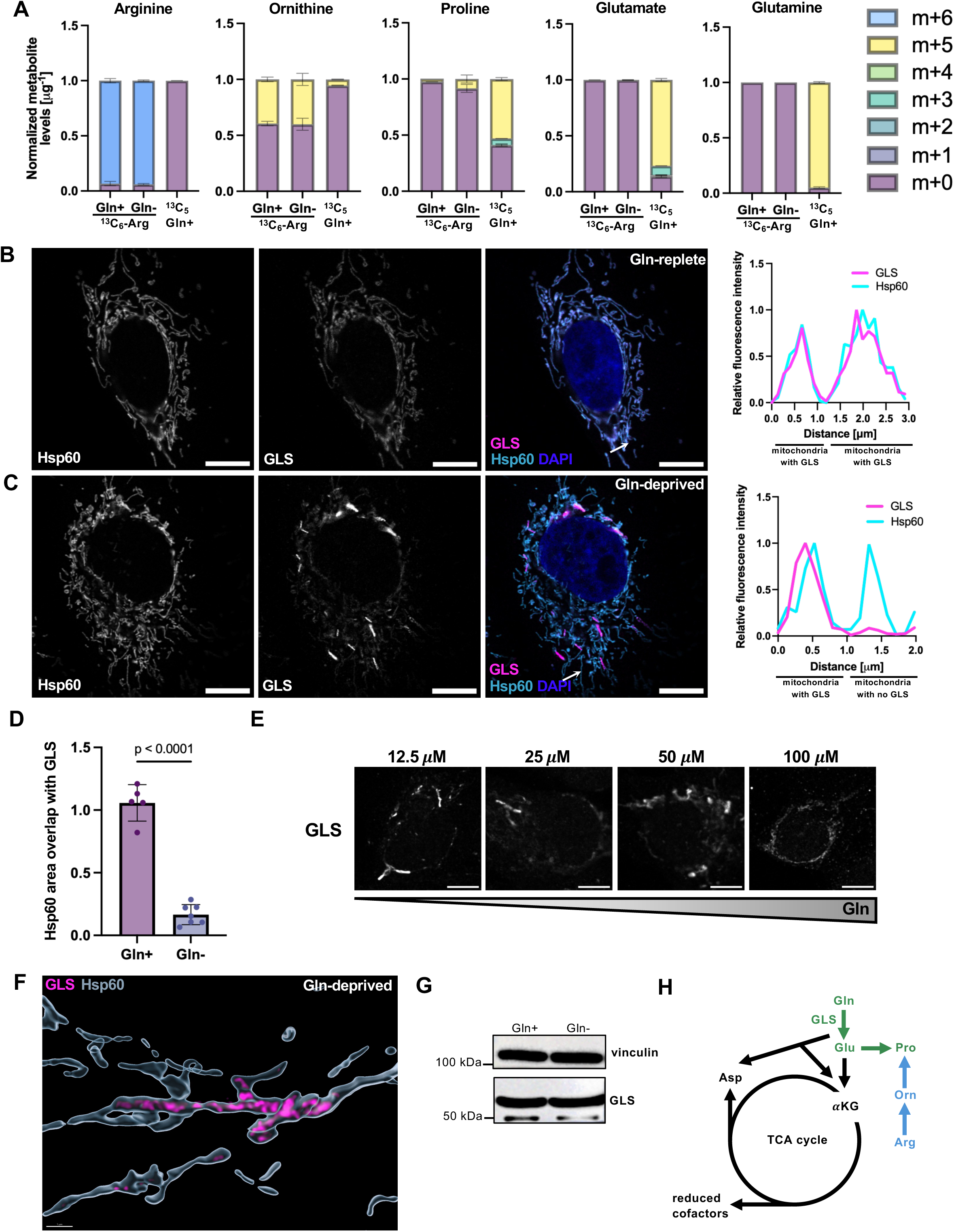
Glutamine deprivation induces GLS redistribution into a distinct mitochondrial subpopulation. **(A)** Fraction of isotopologues for indicated metabolites obtained through tracing with ^13^C_6_-arginine in glutamine-replete and glutamine-deprived conditions, and ^13^C_5_-glutamine in glutamine-replete conditions. Data from *N=*3 independent replicates. **(B)** Representative immunofluorescence image and line scan of GLS distribution within the mitochondrial network under glutamine-replete conditions (4 mM) in Hs 578T cells. Scale bar 10 *μ*m. **(C)** Representative immunofluorescence image and line scan of GLS distribution within the mitochondrial network under glutamine-deprived conditions (25 *μ*M) in Hs 578T cells. Scale bar 10 *μ*m. **(D)** Quantified area overlaps between GLS and Hsp60 as a pan-mitochondrial marker. Statistical significance determined using the two-tailed Student’s t-test. **(E)** Representative micrographs of titration of exogenous glutamine and its impact on GLS redistribution in Hs 578T cells. **(F)** Super-resolution Airyscan images of GLS-positive foci and elongated structures within mitochondria (Hsp60 marker) under glutamine deprivation (3D reconstructed signal) in Hs 578T cells. Scale bar 1 *μ*m. **(G)** Western blot showing GLS expression levels under glutamine-replete and glutamine-deprived conditions in MDA-MB-231 cells. **(H)** Schematic of downstream metabolic fates of glutamine in glutamine-dependent cancer cells, with proline synthesis from glutamine highlighted in green, and the inactive proline synthesis from arginine highlighted in blue.

**Supplementary Figure 2.**
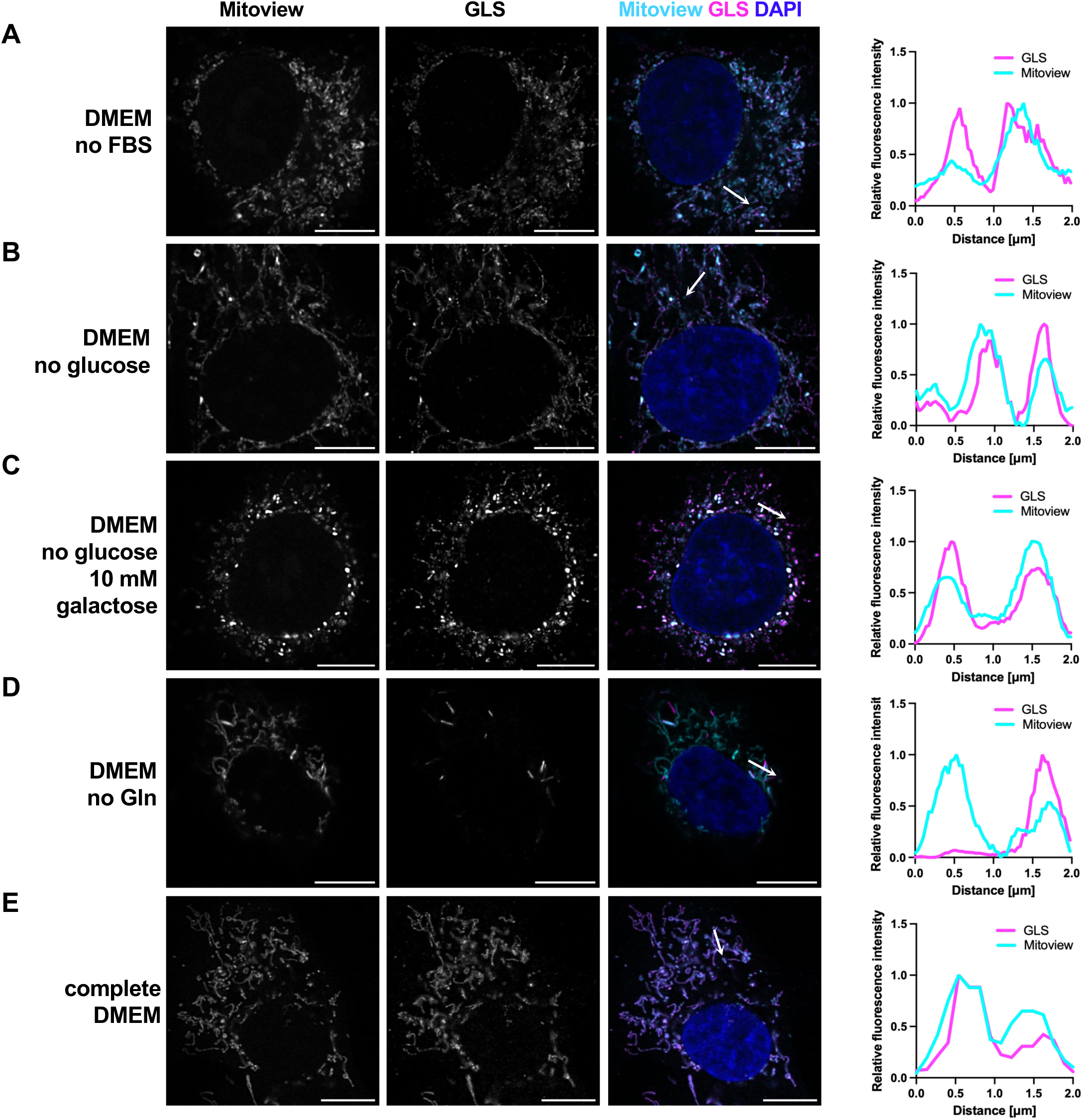
GLS rod formation is selectively induced by glutamine deprivation. **(A)** Representative immunofluorescence image and line scan showing pan-mitochondrial distribution of GLS in the mitochondrial network of Hs 578T cells in FBS-deprived DMEM. Scale bar 10 *μ*m. **(B)** Representative immunofluorescence image and line scan showing pan-mitochondrial distribution of GLS in the mitochondrial network of Hs 578T cells in DMEM without glucose. Scale bar 10 *μ*m. **(C)** Representative immunofluorescence image and line scan showing pan-mitochondrial distribution of GLS in the mitochondrial network of Hs 578T cells in DMEM without glucose supplemented with galactose. Scale bar 10 *μ*m. **(D)** Representative immunofluorescence image and line scan showing cellular rods of GLS forming in glutamine-deprived DMEM. Scale bar 10 *μ*m. **(E)** Representative immunofluorescence image and line scan showing pan-mitochondrial distribution of GLS in the mitochondrial network of Hs 578T cells in complete DMEM. Scale bar 10 *μ*m.

**Supplementary Figure 3.**
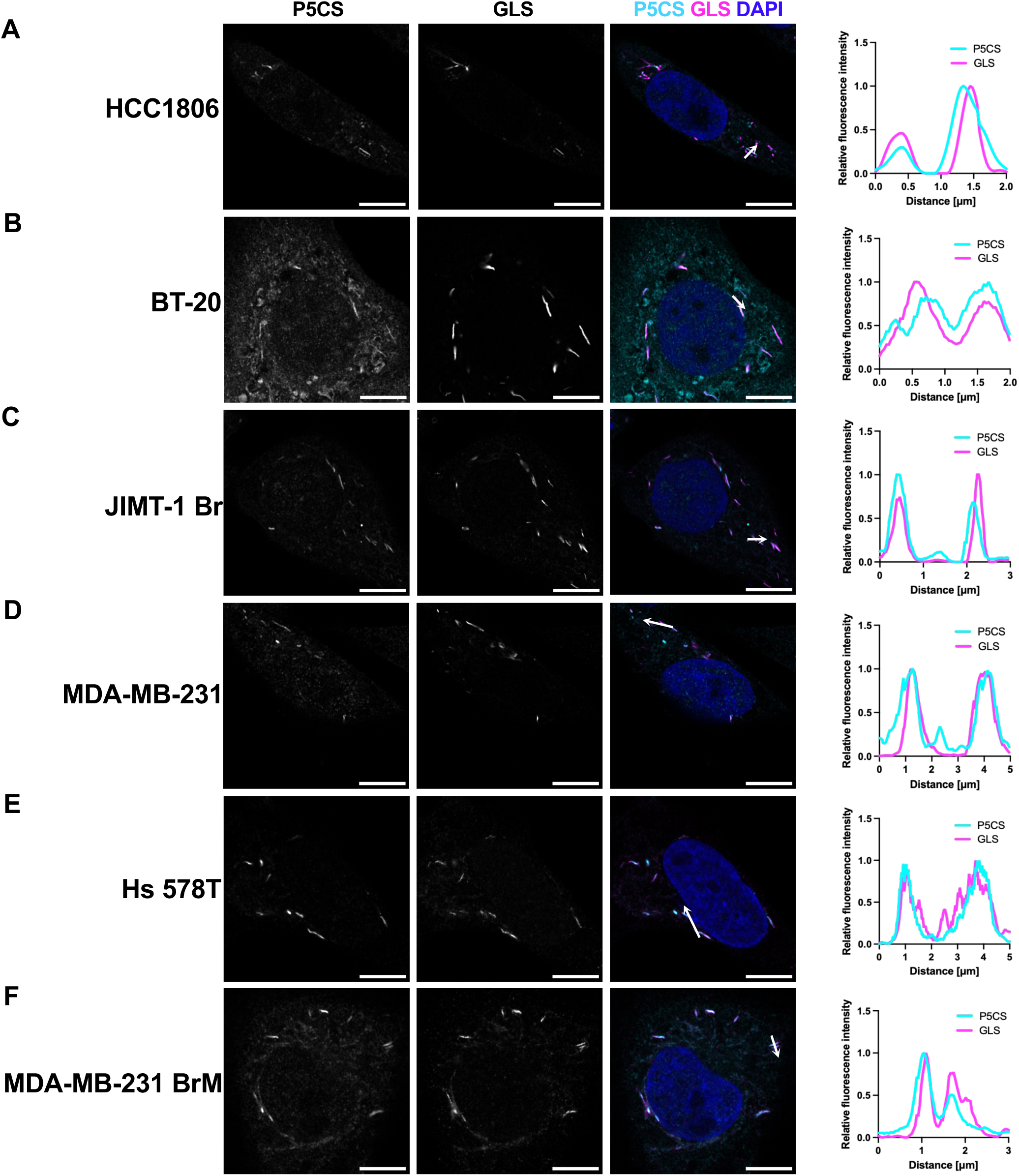
Co-segregation of GLS and P5CS is conserved across glutamine-dependent breast cancer cell lines. **(A)** Representative immunofluorescence image and line scan of P5CS and GLS co-enrichment in mitochondrial rods under glutamine deprivation in HCC1806 cells. Scale bar 10 *μ*m. **(B)** Representative immunofluorescence image and line scan of P5CS and GLS co-enrichment in mitochondrial rods under glutamine deprivation in BT-20 cells. Scale bar 10 *μ*m. **(C)** Representative immunofluorescence image and line scan of P5CS and GLS co-enrichment in mitochondrial rods under glutamine deprivation in JIMT-1 Br cells. Scale bar 10 *μ*m. **(D)** Representative immunofluorescence image and line scan of P5CS and GLS co-enrichment in mitochondrial rods under glutamine deprivation in MDA-MB-231 cells. Scale bar 10 *μ*m. **(E)** Representative immunofluorescence image and line scan of P5CS and GLS co-enrichment in mitochondrial rods under glutamine deprivation in Hs 578T cells. Scale bar 10 *μ*m. **(F)** Representative immunofluorescence image and line scan of P5CS and GLS co-enrichment in mitochondrial rods under glutamine deprivation in MDA-MB-231 brain-metastatic breast cancer cells. Scale bar 10 *μ*m.

**Supplementary Figure 4.**
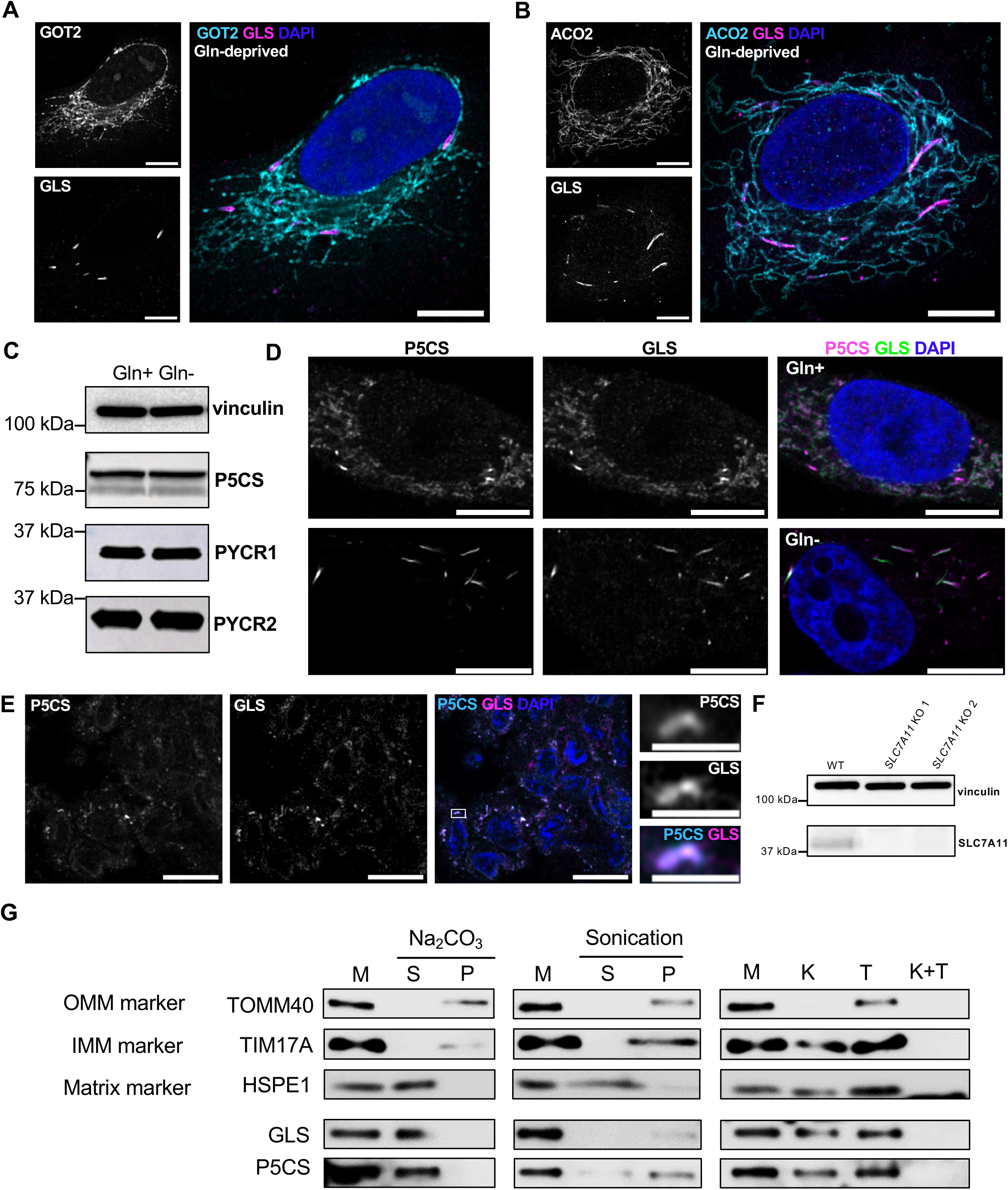
GLS and P5CS selectively co-segregate and are enriched at the inner mitochondrial membrane. **(A)** Representative immunofluorescence image of the pan-mitochondrial distribution of aspartate aminotransferase 2 (GOT2), a mitochondrial matrix enzyme, under glutamine deprivation. Scale bar 10 *μ*m. **(B)** Representative immunofluorescence image of the pan-mitochondrial distribution of aconitase 2 (ACO2), a mitochondrial matrix enzyme, under glutamine deprivation. Scale bar 10 *μ*m. **(C)** Western blot showing P5CS, PYCR1, and PYCR2 expression levels under glutamine-replete and glutamine-deprived conditions in MDA-MB-231 cells. **(D)** Representative immunofluorescence image of the pan-mitochondrial distribution of P5CS and GLS in MDA-MB-231 cells, knockouts for endogenous *GLS* and re-expressing GAC-GFP, the most abundant splice variant of GLS. Upper panel: glutamine-replete conditions, lower panel: glutamine-deprived conditions. Scale bar 10 *μ*m. **(E)** Representative immunofluorescence image of a human breast tumor slice, co-stained for GLS and P5CS, showing rod morphology for both enzymes. Scale bar 10 *μ*m. Insets show an individual rod, scale bar 2 *μ*m. **(F)** Western blot showing SLC7A11 expression levels in wild-type and knockout cells, MDA-MB-231 cell line. **(G)** Western blot of mitochondrial subfractions from MDA-MB-231 cells. TOMM40 used as an outer mitochondrial membrane marker, TIM17A used as an inner mitochondrial membrane marker, HSPE1 used as a matrix marker. In all panels ‘M’ denotes whole mitochondrial input. Left panel: treatment with sodium carbonate decouples membrane-associated proteins and soluble proteins (supernatant, S) from membrane-integral proteins (P). Middle panel: sonication disrupts membranes and separates membranes with membrane-associated proteins (pellet, P) from soluble proteins (supernatant, S). Right panel: Treatment with protease K (K) disrupts outer membrane proteins. Treatment with Triton X-100 (T) solubilises all proteins. Combined treatment degrades all proteins (K+T).

**Supplementary Figure 5.**
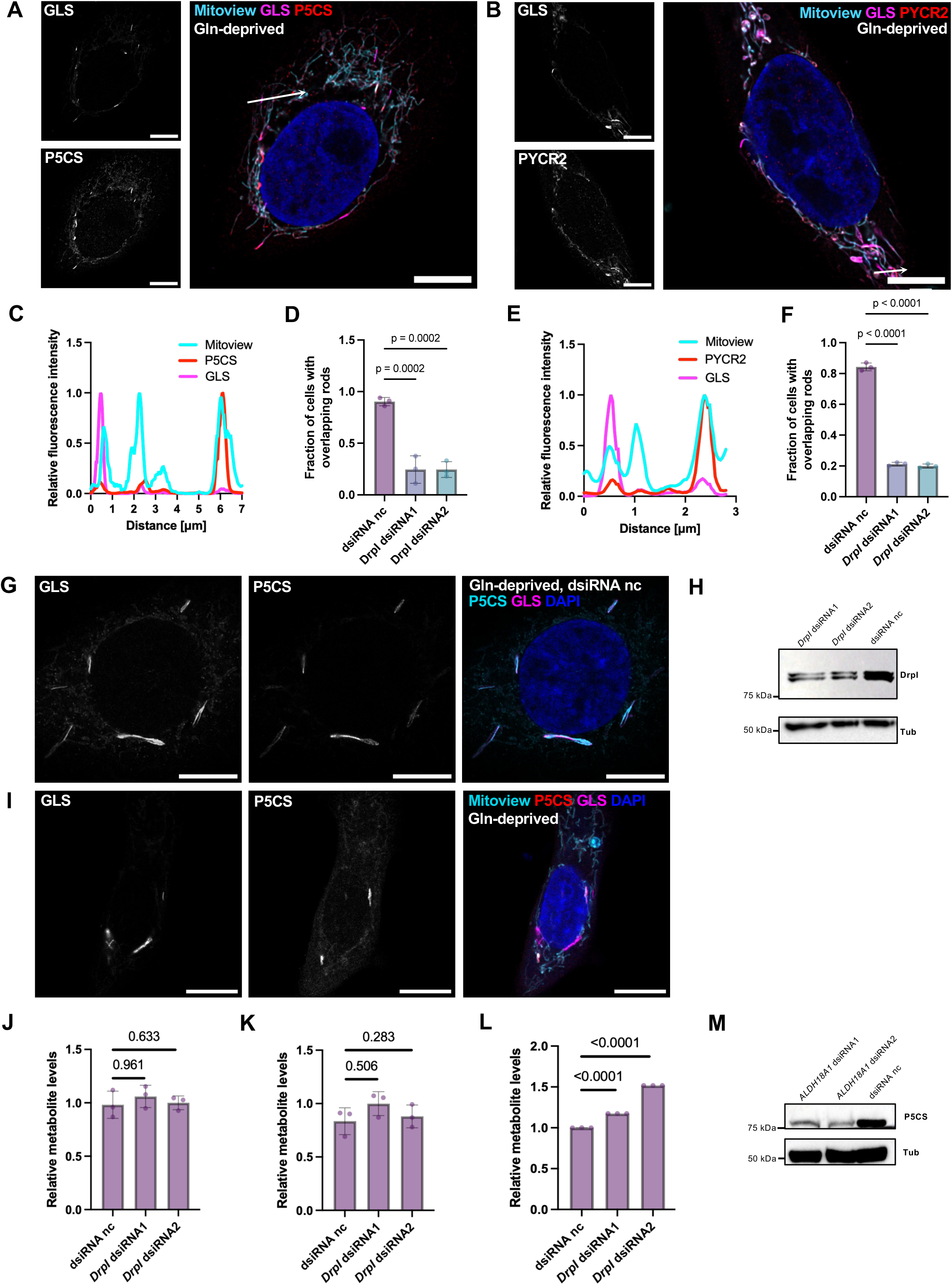
DRP1 depletion disrupts coordinated enzyme segregation without preventing rod formation. **(A)** Representative micrograph and inset of disrupted coordination between GLS and P5CS under glutamine deprivation in *DNM1L* (DRP1) knockdown in Hs 578T cells. Scale bar 10 *μ*m. **(B)** Representative micrograph and inset of disrupted coordination between GLS and PYCR2 under glutamine deprivation in *DNM1L* (DRP1) knockdown in Hs 578T cells. Scale bar 10 *μ*m. **(C)** Line scan for P5CS and GLS in *DNM1L* (DRP1) knockdown in Hs 578T cells. **(D)** Percent cells with coordinated rod segregation of P5CS and GLS into mitochondrial rods in *DNM1L* (DRP1) knockdown Hs 578T cells. Data from *N=*3 independent replicates, statistical significance determined using a one-way ANOVA Tukey’s multiple comparisons test. **(E)** Line scan for PYCR2 and GLS in *DNM1L* (DRP1) knockdown Hs 578T cells. **(F)** Percent cells with coordinated rod segregation of PYCR1 and GLS in *DRPI* knockdown Hs 578T cells. Data from *N=*3 independent replicates, statistical significance determined using a one-way ANOVA with Tukey’s multiple comparisons test. **(G)** Representative micrograph of rod formation in glutamine deprived Hs 578T cells treated with a control dsiRNA. Scale bar 10 *μ*m. **(H)** Western blot showing knockdowns of *DNM1L* (DRP1) in Hs 578T cell line. **(I)** Representative micrograph of co-segregation disruption between GLS and P5CS in glutamine-deprived Hs 578T cells treated with 10 *μ*M DRP1i27 for 24 h. Scale bar 10 *μ*m. **(J)** Relative metabolite levels for arginine in *DNM1L* (DRP1) knockdown cells subject to a 24 h glutamine deprivation with ^13^C_5_-glutamine traced after a 1 h pulse at 1 mM. No label incorporation was detected. Data normalised to levels in control dsiRNA group. Data from *N=*3 independent replicates, statistical significance determined on log2-transformed data using a one-way ANOVA with Tukey’s multiple comparisons test. **(K)** Relative metabolite levels for ornithine in *DNM1L* (DRP1) knockdown cells subject to a 24 h glutamine deprivation with ^13^C_5_-glutamine traced after a 1 h pulse at 1 mM. No label incorporation was detected. Data normalised to levels in control dsiRNA group. Data from *N=*3 independent replicates, statistical significance determined on log2-transformed data using a one-way ANOVA with Tukey’s multiple comparisons test. **(L)** Relative metabolite levels for *m+5* glutamine in *DNM1L* (DRP1) knockdown cells subject to a 24 h glutamine deprivation with ^13^C_5_-glutamine traced after a 1 h pulse at 1 mM. Data normalised to levels in control dsiRNA group. Data from *N=*3 independent replicates, statistical significance determined on log2-transformed data using a one-way ANOVA with Tukey’s multiple comparisons test. **(M)** Western blot showing knockdowns of *ALDH18A1* (protein product P5CS) in Hs 578T cells.

**Movie S1.**

Cellular rod formation of GLS and P5CS and relative tri-dimensional reconstructions of the signal.

**Movie S2.**

Cellular rod formation of GLS and PYCR1 and relative tri-dimensional reconstructions of the signal.

**Movie S3.**

Cellular rod formation of GLS and PYCR2 and relative tri-dimensional reconstructions of the signal.

## References

1. De Visser, K. E. & Joyce, J. A. The evolving tumor microenvironment: From cancer initiation to metastatic outgrowth. Cancer Cell 41, 374–403 (2023).

2. DeBerardinis, R. J. & Chandel, N. S. Fundamentals of cancer metabolism. Sci. Adv. 2, e1600200 (2016).

3. Hanahan, D. Hallmarks of cancer—Then and now, and beyond. Cell 189, 2254–2277 (2026).

4. Cluntun, A. A., Lukey, M. J., Cerione, R. A. & Locasale, J. W. Glutamine Metabolism in Cancer: Understanding the Heterogeneity. Trends in Cancer 3, 169–180 (2017).

5. Yoo, H. C., Yu, Y. C., Sung, Y. & Han, J. M. Glutamine reliance in cell metabolism. Exp Mol Med 52, 1496–1516 (2020).

6. Sweetlove, L. J. & Fernie, A. R. The role of dynamic enzyme assemblies and substrate channelling in metabolic regulation. Nat Commun 9, 2136 (2018).

7. Pareek, V., Sha, Z., He, J., Wingreen, N. S. & Benkovic, S. J. Metabolic channeling: predictions, deductions, and evidence. Molecular Cell 81, 3775–3785 (2021).

8. Zhang, Y. & Fernie, A. R. Metabolons, enzyme–enzyme assemblies that mediate substrate channeling, and their roles in plant metabolism. Plant Communications 2, 100081 (2021).

9. Ryu, K. W. et al. Cellular ATP demand creates metabolically distinct subpopulations of mitochondria. Nature 635, 746–754 (2024).

10. Yang, C. et al. De novo pyrimidine biosynthetic complexes support cancer cell proliferation and ferroptosis defence. Nat Cell Biol 25, 836–847 (2023).

11. Fuller, G. G. et al. RNA promotes phase separation of glycolysis enzymes into yeast G bodies in hypoxia. eLife 9, e48480 (2020).

12. French, J. B. et al. Spatial colocalization and functional link of purinosomes with mitochondria. Science 351, 733–737 (2016).

13. Park, C. K. & Horton, N. C. Structures, functions, and mechanisms of filament forming enzymes: a renaissance of enzyme filamentation. Biophys Rev 11, 927–994 (2019).

14. Narayanaswamy, R., et al. Widespread reorganization of metabolic enzymes into reversible assemblies upon nutrient starvation. Proc. Natl. Acad. Sci. U.S.A. 106, 10147–10152 (2009).

15. Lynch, E. M., Kollman, J. M. & Webb, B. A. Filament formation by metabolic enzymes— A new twist on regulation. Current Opinion in Cell Biology 66, 28–33 (2020).

16. Adamoski, D. et al. Molecular mechanism of glutaminase activation through filamentation and the role of filaments in mitophagy protection. Nat Struct Mol Biol 30, 1902–1912 (2023).

17. Lynch, E. M. et al. Structural basis for allosteric regulation of human phosphofructokinase-1. Nat Commun 15, 7323 (2024).

18. Lynch, E. M. et al. Human CTP synthase filament structure reveals the active enzyme conformation. Nat Struct Mol Biol 24, 507–514 (2017).

19. Ferreira, A. P. S. et al. Active Glutaminase C Self-assembles into a Supratetrameric Oligomer That Can Be Disrupted by an Allosteric Inhibitor. Journal of Biological Chemistry 288, 28009–28020 (2013).

20. Jiang, B. et al. Filamentous GLS1 promotes ROS-induced apoptosis upon glutamine deprivation via insufficient asparagine synthesis. Molecular Cell 82, 1821–1835.e6 (2022).

21. Zhao, W., et al. HIF2α inhibits glutaminase clustering in mitochondria to sustain growth of clear cell renal cell carcinoma. JCI Insight 10, e182711 (2025).

22. Gunn, K. H., et al. The structure of helical lipoprotein lipase reveals an unexpected twist in lipase storage. Proc. Natl. Acad. Sci. U.S.A. 117, 10254–10264 (2020).

23. Gunn, K. H. et al. Cryogenic electron tomography reveals helical organization of lipoprotein lipase in storage vesicles. Sci. Adv. 11, eadx8711 (2025).

24. Cassago, A., et al. Mitochondrial localization and structure-based phosphate activation mechanism of Glutaminase C with implications for cancer metabolism. Proc. Natl. Acad. Sci. U.S.A. 109, 1092–1097 (2012).

25. Zhong, J. et al. Structural basis of dynamic P5CS filaments. eLife 11, e76107 (2022).

26. Feng, S., Aplin, C., Nguyen, T.-T. T., Milano, S. K. & Cerione, R. A. Filament formation drives catalysis by glutaminase enzymes important in cancer progression. Nat Commun 15, 1971 (2024).

27. Koppula, P., Zhang, Y., Zhuang, L. & Gan, B. Amino acid transporter SLC7A11/xCT at the crossroads of regulating redox homeostasis and nutrient dependency of cancer. Cancer Commun 38, 12 (2018).

28. Timmerman, L. A. et al. Glutamine Sensitivity Analysis Identifies the xCT Antiporter as a Common Triple-Negative Breast Tumor Therapeutic Target. Cancer Cell 24, 450–465 (2013).

29. Shin, C.-S. et al. The glutamate/cystine xCT antiporter antagonizes glutamine metabolism and reduces nutrient flexibility. Nat Commun 8, 15074 (2017).

30. Muir, A. et al. Environmental cystine drives glutamine anaplerosis and sensitizes cancer cells to glutaminase inhibition. eLife 6, e27713 (2017).

31. Qiu, Y. et al. The unique catalytic properties of PSAT1 mediate metabolic adaptation to glutamine blockade. Nat Metab 6, 1529–1548 (2024).

32. Muir, A. & Vander Heiden, M. G. The nutrient environment affects therapy. Science 360, 962–963 (2018).

33. Petrovska, I. et al. Filament formation by metabolic enzymes is a specific adaptation to an advanced state of cellular starvation. eLife 3, e02409 (2014).

34. McCorvie, T. J. et al. Architecture and regulation of filamentous human cystathionine beta-synthase. Nat Commun 15, 2931 (2024).

35. Fung, T. S., et al. Mitochondrial subpopulations promote cellular responses to changes in metabolic demand and signal transduction. Proc. Natl. Acad. Sci. U.S.A. 123, e2613074123 (2026).

36. Guo, C.-J. et al. Dynamic Arabidopsis P5CS filament facilitates substrate channelling. Nat. Plants 10, 880–889 (2024).

37. Christensen, E. M. et al. Resolving the cofactor-binding site in the proline biosynthetic enzyme human pyrroline-5-carboxylate reductase 1. Journal of Biological Chemistry 292, 7233– 7243 (2017).

38. Bogner, A. N., Stiers, K. M. & Tanner, J. J. Structure, biochemistry, and gene expression patterns of the proline biosynthetic enzyme pyrroline-5-carboxylate reductase (PYCR), an emerging cancer therapy target. Amino Acids 53, 1817–1834 (2021).

39. Yang, Z. et al. Pyrroline-5-carboxylate synthase senses cellular stress and modulates metabolism by regulating mitochondrial respiration. Cell Death Differ 28, 303–319 (2021).

40. Matés, J. M. et al. Glutamine homeostasis and mitochondrial dynamics. The International Journal of Biochemistry & Cell Biology 41, 2051–2061 (2009).

41. Schwörer, S. et al. Proline biosynthesis is a vent for TGFβ-induced mitochondrial redox stress. EMBO J 39, EMBJ2019103334 (2020).

42. Zhu, J. et al. Mitochondrial NADP(H) generation is essential for proline biosynthesis. Science 372, 968–972 (2021).

43. Burke, L. et al. The Janus-like role of proline metabolism in cancer. Cell Death Discov. 6, 104 (2020).

44. Santiappillai, N. T. et al. Macrophage-supported metabolic adaptation drives tumor survival under nutrient stress. Sci. Adv. 12, eaei2831 (2026).

45. Sullivan, L. B. et al. Supporting Aspartate Biosynthesis Is an Essential Function of Respiration in Proliferating Cells. Cell 162, 552–563 (2015).

46. Linder, S. J. et al. Inhibition of the proline metabolism rate-limiting enzyme P5CS allows proliferation of glutamine-restricted cancer cells. Nat Metab 5, 2131–2147 (2023).

47. Zhou, D. et al. Protocol for mitochondrial isolation and sub-cellular localization assay for mitochondrial proteins. STAR Protocols 4, 102088 (2023).

48. Garg, A., Shang, R., Cvetanovic, T., Lai, E. C. & Joshua-Tor, L. The structural landscape of Microprocessor-mediated processing of pri-let-7 miRNAs. Molecular Cell 84, 4175–4190.e6 (2024).

